# Spatial transcriptomics unveils the *in situ* cellular and molecular hallmarks of the lung in fatal COVID-19

**DOI:** 10.1101/2024.07.03.601404

**Authors:** Carlos A. Garcia-Prieto, Eva Musulen, Veronica Davalos, Gerardo Ferrer, Daniela Grases, Eduard Porta, Belén Pérez-Miés, Tamara Caniego-Casas, José Palacios, Xavier Saenz-Sardà, Elisabet Englund, Manel Esteller

## Abstract

Severe Coronavirus disease 2019 (COVID-19) induces heterogeneous and progressive diffuse alveolar damage (DAD) highly disrupting lung tissue architecture and homeostasis, hampering disease management leading to fatal outcomes. Characterizing DAD pathophysiology across disease progression is of ultimate importance to better understand the molecular and cellular features driving different DAD patterns and to optimize treatment strategies. To contextualize the interplay between cell types and assess their distribution, spatial transcriptomics (ST) techniques have emerged, allowing unprecedented resolution to investigate spatial architecture of tissues. To this end, post-mortem lung tissue provides valuable insights into cellular composition and their spatial relationships at the time of death. Here, we have leveraged VisumST technology in post-mortem COVID-19 induced acute and proliferative DAD lungs including control samples with normal morphological appearance, to unravel the immunopathological mechanisms underlying DAD, providing novel insights into cellular and molecular communication events driving DAD progression in fatal COVID-19. We report a progressive loss of endothelial cell types, pneumocytes type I and natural killer cells coupled with a continuous increase of myeloid and stromal cells, mostly peribronchial fibroblasts, over disease progression. Spatial organization analysis identified variable cellular compartments, ranging from major compartments defined by cell type lineages in control lungs to increased and more specific compartmentalization including immune-specific clusters across DAD spectrum. Importantly, spatially informed ligand-receptor interaction (LRI) analysis revealed an intercellular communication signature defining COVID-19 induced DAD lungs. Transcription factor (TF) activity enrichment analysis identified TGF-B pathway as DAD driver, highlighting SMAD3 and SMAD7 TFs activity role during lung fibrosis. Integration of deregulated LRIs and TFs activity allowed us to propose a downstream intracellular signaling pathway in peribronchial fibroblasts, suggesting potential novel therapeutic targets. Finally, spatio-temporal trajectories analysis provided insights into the alveolar epithelium regeneration program, characterizing markers of pneumocytes type II differentiation towards pneumocytes type I. In conclusion, we provide a spatial characterization of lung tissue architecture upon COVID-19 induced DAD progression, identifying molecular and cellular hallmarks that may help optimize treatment and patient management.

## Introduction

Infection by the severe acute respiratory syndrome coronavirus 2 (SARS-CoV-2) virus caused a worldwide pandemic of the derived coronavirus disease 2019, COVID-19. At the time of the writing (June 28^th^, 2023) beyond 775 million confirmed cases and more than 7 million deaths (https://data.who.int/dashboards/covid19/) had been reported. COVID-19 has a wide range of symptomatology, but most affected individuals exhibit mild clinical manifestations or are asymptomatic (Coronaviridae Study Group, 2020; Wu et al., 2020). Importantly, respiratory failure linked to lung damage and acute respiratory distress syndrome (ARDS) is the most common cause of death in COVID-19 patients (Milross et al., 2022; Bridges et al., 2022). Mechanical ventilation to compensate for respiratory failure is usually required for these high-risk cases (Berlin et al., 2020; Marini et al., 2020). Most importantly, for the current clinical situation with the advancement of COVID-19 vaccines, the maintenance of lung lesions in a subgroup of COVID-19 patients could also be associated to prolonged clinical manifestation (Adeloye et al., 2021; Davis, et al., 2023). Upon SARS-CoV-2 infection in the severe cases, the pathological change is defined by the presence of diffuse alveolar damage (DAD) (Erjefält et al., 2022) in the lung that initiates with a first acute stage of early intra-alveolar epithelial lesions, interstitial inflammation and oedema, followed by, the proliferative stage with a final appearance of pneumocyte hyperplasia and fibroblast proliferation (Milross et al., 2022; Bridges et al., 2022).

Despite the principal contribution of respiratory failure to lethal COVID-19, the molecular context of lung damage provoked by SARS-CoV-2 infection is not fully established. Among the multiple multiomics layers that could be interrogated in the disorder, most studies have only addressed the bulk transcription landscape of the disease (Blanco-Melo et al., 2020; Pinto et al., 2020; D’Agnillo et al., 2021) or a particularly isolated lineage such as immune cells (Liao et al., 2020; Wilk et al, 2020). A more granulated view of the lung affected by severe COVID- 19 can be gained by the characterization of single-cell gene expression profiles as it has also been recently reported (Mels et al., 2021; Delorey et al., 2021; Sikkema et al., 2023). However, a comprehensive and unbiased transcriptional profiling of the lung in these severe cases of COVID-19 have not been properly available since all previous approaches destroyed the rich and relevant anatomical structure of the lung that could be crucial in understanding its pathobiology. Even if in the above approaches there is purification of particularly important cell types, the global determination of cell-cell interactions in the space of the host tissue is lost. In this regard, only a limited number of studies have analyzed a spatial component in the lung of COVID-19 cases, using high-parameter imaging mass cytometry for a discreet set of targeted proteins (Rendeiro et al., 2021) or restricted to small sublocations termed regions of interest (ROI) (Desai et al., 2020; Margaroli et al., 2021; Park et al., 2022; Milross et al., 2024).

Importantly, larger pathological scrutiny of spatial transcriptomics in COVID-19 patients has only been performed recently and to the best of our knowledge, once (Mothes et al., 2023).

To overcome these issues, we leveraged the recently developed Visium spatial transcriptomics (VisiumST) technology in a cohort of lungs with normal histology and those that underwent DAD upon the course of fatal COVID-19. To further untangle the precise and topographically located gene expression changes underlying the disease, both acute and proliferative stages of DAD were studied using the single-cell RNA expression data of a human cell atlas of the lung to annotate cell types (Sikkema et al., 2023). These exhaustive analyses provided the constellation of shifts in the forty-five cell types and fates annotated that occur upon severe COVID-19, but also the perturbations in cell-to-cell communication events. The unveiled anatomical specific information of the altered cell types, their spatial relationships and their corresponding molecular pathways could be extremely valuable to design more targeted-based pharmacological treatments to prevent the progression of the disease in severe COVID- 19 patients.

## Results

### Identification of cell types and cellular compartments in the lung through COVID-19 associated DAD progression

To assess how fatal COVID-19 affected cell type composition, cell-cell communication, and global expression patterns across DAD progression in the lung of patients that died by the disease, we followed the study design shown in **Figure 1A**. We first retrieved twenty-three formalin-fixed paraffin-embedded (FFPE) post-mortem lung tissue samples obtained from nineteen patients with DAD, corresponding to seven cases of acute DAD stage and twelve proliferative DAD stages classified as previously described (Pérez-Mies et al., 2022), and four lung samples from control lungs with normal morphological appearance without clinical evidence of SARS-CoV-2 infection. The clinicopathological characteristics of the studies samples are described in **Table 1**. We then analyzed the spatial transcriptomics patterns of the described samples using tissue spots on a microarray slide with arrayed oligonucleotides to capture spatial gene expression information (Ståhl et al., 2016). Reverse transcription was performed on the intact tissue, and the resulting cDNA was coupled to the oligonucleotides on the slide before tissue lysis, before final generation of next generation sequencing (NGS) libraries (Ståhl et al., 2016). This technology was later adapted by 10x Genomics as ‘10x Visium’ (VisiumST), with increased resolution of 55 μm (10X Genomics, 2019), as we have herein used as previously described (Rao et al., 2021). In total, 91,068 tissue spots were studied after quality control (QC) and preprocessing (**Methods**), selecting the most informative genes as previously described (Sikkema et al., 2023) (**Methods**). The herein used spatial transcriptomics technology does not achieve complete single-cell level approach. To assess the spatial organization of cell types across tissue slides, we used the integrated Human Lung Cell Atlas (HLCA), that combines 49 single-cell RNA datasets spanning 2.4 million cells from 486 individuals (Sikkema et al., 2023), as a reference to deconvolute the main cell types present in each spot by applying the validated cell2location pipeline (Kleshchevnikov et al., 2022; Madissoon et al., 2023). Uniform Manifold Approximation and Projection (UMAP) using the integrated global spatial transcriptome data showing the spots colored by our three groups of lung samples (control, acute DAD, and proliferative DAD) was able to distinguish these three entities (**Figure 1B**). Both the entire set of cases and illustrative examples are shown in **Figure 1B**.

**Figure 1:**
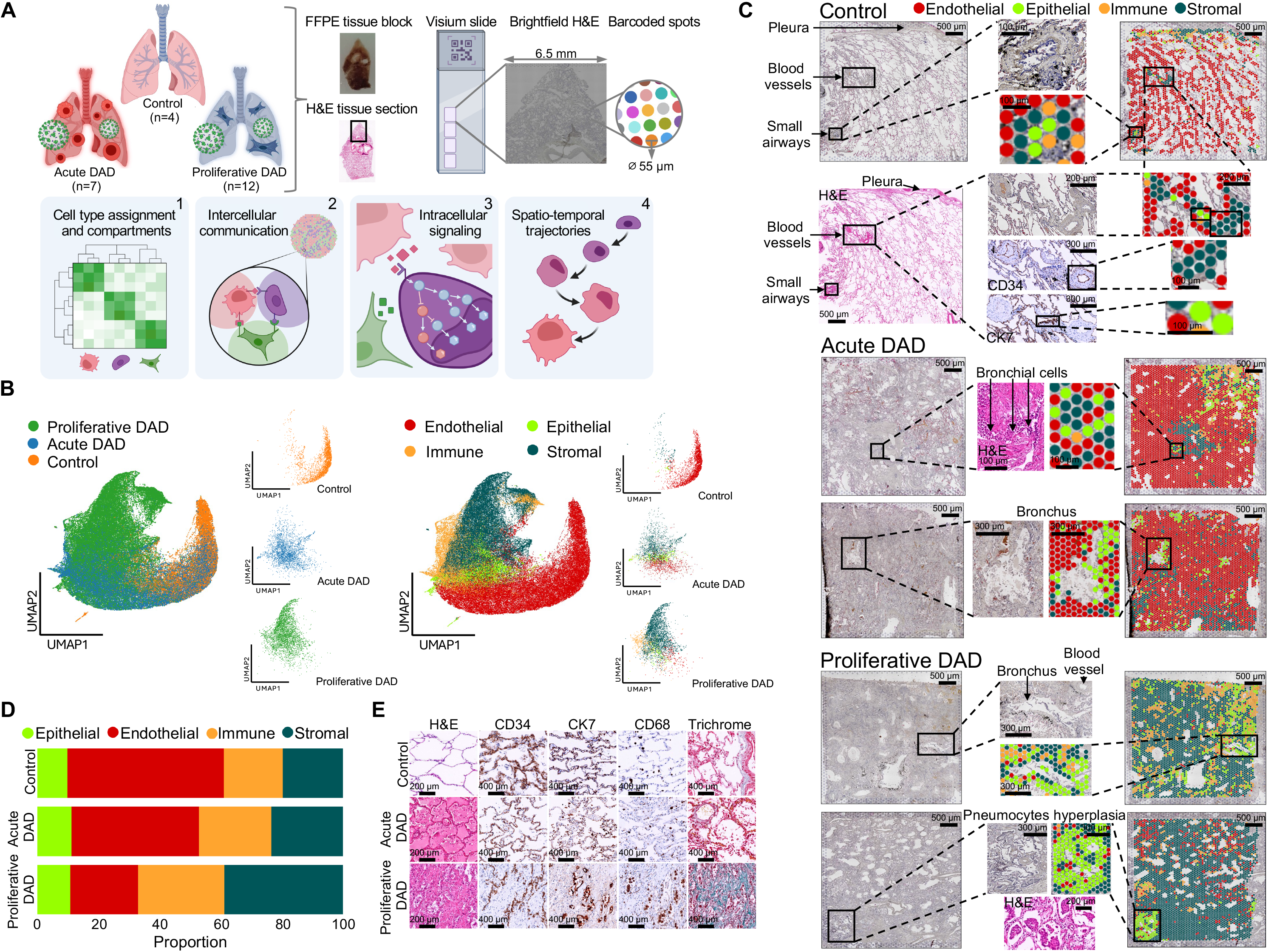
Study design and spatial transcriptomics profiling of fatal COVID-19. **A)** Overview of the study design, including sample processing workflow and spatial transcriptomics data analysis pipeline. **B)** UMAP representing sample integration and spot-wise most abundant cell type assignment. **C)** Mapping cell type deconvolution results on top of VisiumST brightfield images stained with H&E across disease progression. Illustrative examples of lung structures are shown matching cell types to expected structures. **D)** Bar plot showing cell type lineage abundance per condition. **E)** Immunohistochemistry staining with H&E and specific markers of the four main cell type lineages: CK7 (alveolar epithelial cells), CD34 (endothelial cells), CD68 (alveolar macrophages) and trichrome (fibrosis).

**Table 1.** Clinicopathological characteristics of the studied COVID-19 patients and control group.

| Characteristics | COVID-19 cohort<br>(N = 19)<sup>a</sup> | Control cohort<br>(N = 4)<sup>b</sup> | p-value* |
| --- | --- | --- | --- |
| Gender - Frequency (%) |  |  |  |
| Female | 5 (26.3) | 2 (50.0) | p=0.557 |
| Male | 14 (73.7) | 2 (50.0) |  |
| Age (years) - Median [range] | 68.0 [52 - 91] | 65.5 [39 - 72] | p=0.409 |
| Underlying conditions - Frequency (%) |  |  |  |
| Smoking | 5 (38.5) | 1 (25.0) | p=1.000 |
| Hypertension | 12 (63.2) | 2 (50.0) | p=1.000 |
| Diabetes mellitus | 3 (15.8) | 1 (25.0) | p=1.000 |
| Obesity | 9 (52.9) | 1 (25.0) | p=0.586 |
| Respiratory disease<sup>c</sup> | 5 (26.3) | 0 (0.00) | p=0.539 |
| Cardiac disease<sup>d</sup> | 13 (68.4) | 0 (0.00) | p=0.024 |
| Chronic kidney disease | 2 (10.5) | 0 (0.00) | p=1.000 |
| Chronic neurological or neuromuscular disease | 5 (26.3) | 1 (25.0) | p=1.000 |
| Cancer | 7 (36.8) | 2 (50.0) | p=1.000 |
| Immunocompromised state<sup>e</sup> | 3 (15.8) | 2 (50.0) | p=0.194 |
| Number of comorbidities - Frequency (%) |  |  |  |
| 0-2 | 6 (31.6) | 2 (50.0) | p=0.589 |
| 3-6 | 13 (68.4) | 2 (50.0) |  |
| Pulmonary disease - Frequency (%) |  |  |  |
| Acute | 7 (36.8) | NA |  |
| Proliferative | 12 (63.2) | NA |  |
| Cause of death - Frequency (%) |  |  |  |
| Multiorgan failure/Respiratory distress | 14 (73.7) | 0 (0) | p=0.014 |
| Pancreatitis | 2 (10.5) | 0 (0) | p=1.000 |
| Intestinal necrosis | 1 (5.3) | 0 (0) | p=1.000 |
| Cardiopathy<sup>f</sup> | 1 (5.3) | 1 (25.0) | p=0.324 |
| Pyelonephritis | 1 (5.3) | 0 (0) | p=1.000 |
| Cancer dissemination | 0 (0) | 1 (25.0) | p=0.174 |
| Acute pulmonary embolism | 0 (0) | 1 (25.0) | p=0.174 |
| Alive<sup>g</sup> | 0 (0) | 1 (25.0) | NA |

Abbreviations: DAD, diffuse alveolar damage; NA, Not applicable. <sup>a</sup>Inclusion criteria for the COVID-19 cohort: Patients >18 years old, PCR positive for SARS-CoV-2 with complete clinical information of disease history, comorbidities and follow-up, showing clinical pulmonary involvement and COVID-19-related death; <sup>b</sup>Inclusion criteria for the control cohort: >18 years old individuals with complete clinical information about comorbidities, without clinical evidence of SARS-CoV-2 infection, and sudden death due to cardiopathies, except one case died due to cancer dissemination and one post-surgery; <sup>c</sup>Chronic obstructive pulmonary disease or asthma; <sup>d</sup>Coronary artery disease, heart failure or atrial fibrillation; <sup>e</sup>Immunocompromised state due to autoimmune disease or other cause; <sup>f</sup>Acute myocardial infarction, cardiac fibrosis, aortic dissection, hemopericardium or myocarditis; <sup>g</sup>A normal lung biopsy was included in the Control group. \*p-values were calculated using Fisher's exact test or Mann-Whitney test for dichotomous or continuous variables, respectively. p-values under 0.05 represent statistical significant association between co-variables.

Using cell2location, we annotated forty-five cell types in HLCA defined by derived markers (**Methods**) in our lung samples (**Figure S1**). A UMAP visualization of these cell populations according to their lineage (epithelial, stroma, immune and endothelial) in the spatial transcriptomics spots for all the integrated samples and illustrative examples are shown in **Figure 1B**. The mapping of the identified cell types on top of the VisiumST brightfield images stained with Hematoxylin and Eosin (HE) for the described lineages in illustrative control and acute and proliferative DAD cases are shown in **Figure 1C**. Cell types mapped to their expected locations, matching well-described structures, with epithelial cells lining the airway lumen and stromal cells mapping to blood vessel walls, as validated by endothelial (CD34) and epithelial (CK7) markers immunostaining (**Figure 1C**).

Overall, we observed that in control lungs the most abundant lineage corresponded to endothelial cell types, as previously described (Travaglini et al., 2020), and that the acute DAD phase was characterized by a decrease of the endothelial cell types and an increase in immune infiltrates, whereas the proliferative phase was mostly defined by the large proportion of fibrotic tissue, as previously described (Milross et al., 2022; Bridges et al., 2022) and shown in **Figure 1D**. These findings were validated using hematoxylin/eosine (HE) stained sections and specific immunohistochemistry (IHC) markers such as CD34 (endothelial), CD68 (myeloid lineage) and CK7 (epithelial markers) and trichrome staining (fibroblasts) shown in **Figure 1E**.

We then moved to characterize a possible uneven distribution of the identified 45 cell types according to their abundance in the control, acute and proliferative DAD lung groups (**Figure 2A** and **2B**). To ease the analyses interpretation, we ordered these cell populations according to their lineage (epthelial, stromal, immune, and endothelial) (**Figure 2A**). Among the epithelial lineage (10 identified cell types), the most important difference was observed in alveolar type 1 (AT1) cells, comprising between 5-10% of total cells in normal lungs as previously reported (Crapo et al., 1982), that were significantly downregulated in COVID-19 associated proliferative DAD cases in comparison to controls (FDR<0.05) and acute DAD lungs (FDR<0.05) (**Figure 2A and 2B**). The reverse process was observed in alveolar type 2 (AT2) cells that were upregulated in proliferative cases in comparison to acute DAD and control samples (**Figure 2A and 2B**). These results fit with the concept that AT1s are the main cell type responsible for provision of the interface for the blood-gas exchange (a function that it is compromised in COVID-19 patients); whereas AT2 cells function as progenitors that repair the injured alveoli epithelium (Chan and Liu, 2022). Regarding the endothelial lineage (7 identified cell types), we observed that capillary cell types were the most abundant of lung cells (∼30% of total cells), as previously reported (Crapo et al., 1982; Travaglini et al., 2020), and that the studied lung samples underwent a progressive loss from normal lung to acute DAD stage to the final proliferative DAD phase for the abundance of endothelial cells (EC) aerocyte capillary, arterial, general capillary and venous pulmonary (**Figure 2A and 2B**) (for all cases FDR<0.05). These results fully support the mounting evidence linking SARS-CoV-2 infection to multiple endothelial dysfunction (Xu et al., 2023). For the immune lineage (19 identified cell populations), we observed in the myeloid lineage that alveolar macrophages (Mph) CCL3+, and monocyte-derived Mph were significantly overrepresented in COVID-19 associated proliferative DAD cases in comparison to controls (FDR<0.05) and acute DAD lungs (FDR<0.05) (**Figure 2A and 2B**). Interstitial Mph perivascular showed a progressive increase in the evolution of the disease from controls to acute cases (FDR<0.05) and from these to the proliferative samples (FDR<0.05). In the lymphoid lineage, a decrease in CD4 and CD8 T-cells was observed in proliferative COVID-19 patients in comparison to the control group (FDR<0.05) (**Figure 2A and 2B**). Interestingly, Natural Killer (NK) cells experienced a significant decrease from control and acute DAD samples to proliferative DAD cases (FDR<0.05) (**Figure 2A and 2B**). In this regard, these innate effector lymphocytes that respond to acute viral infections have been previously related to COVID-19 severity (Maucourant et al., 2020). Additionally, both Alveolar macrophages and Plasma cells were overrepresented in proliferative DAD lungs compared to control samples (FDR<0.05). Finally, regarding the stromal lineage (9 cell types), the most dramatic change was observed for peribronchial fibroblast with increased numbers in the progression of the disease from control to acute DAD phase, but skyrocketed in proliferative DAD cases (FDR<0.05) (**Figure 2A and 2B**). Not all fibroblast subtypes behaved in a similar manner, including subpleural fibroblasts where we also found an overrepresentation in COVID-19 associated DAD fatal cases compared to the control group and between acute and proliferative stages (FDR<0.05), whereas for alveolar fibroblasts and pericytes we saw a decrease in the proliferative DAD samples compared to control and acute DAD lungs (FDR<0.05), likely associated with the loss of alveoli and endothelial cells characteristic of the proliferative DAD phase of the disease. Lastly, adventitial fibroblasts were overrepresented in proliferative DAD samples compared to controls (FDR<0.05). Illustrative UMAPs for the entire set of cases in the cellular populations of AT1, AT2, EC aerocyte capillary, peribronchial fibroblast, monocyte-derived macrophages and smooth muscle activated stress response cells are shown in **Figure 2C**, showing unbalanced distribution between control, acute and proliferative groups (**Figure 1B**). The mapping of the identified cell types on top of the VisiumST brightfield images stained with HE in illustrative control and COVID-19 associated acute and proliferative DAD cases are shown in **Figure 2B**. As expected, cell types annotated with the finest granularity level mapped to their expected locations, including subpleural fibroblasts located next to the pleura, and multiciliated cells lining small airways lumen and smooth muscle cells mapping to blood vessel walls (**Figure 2B**).

**Figure 2:**
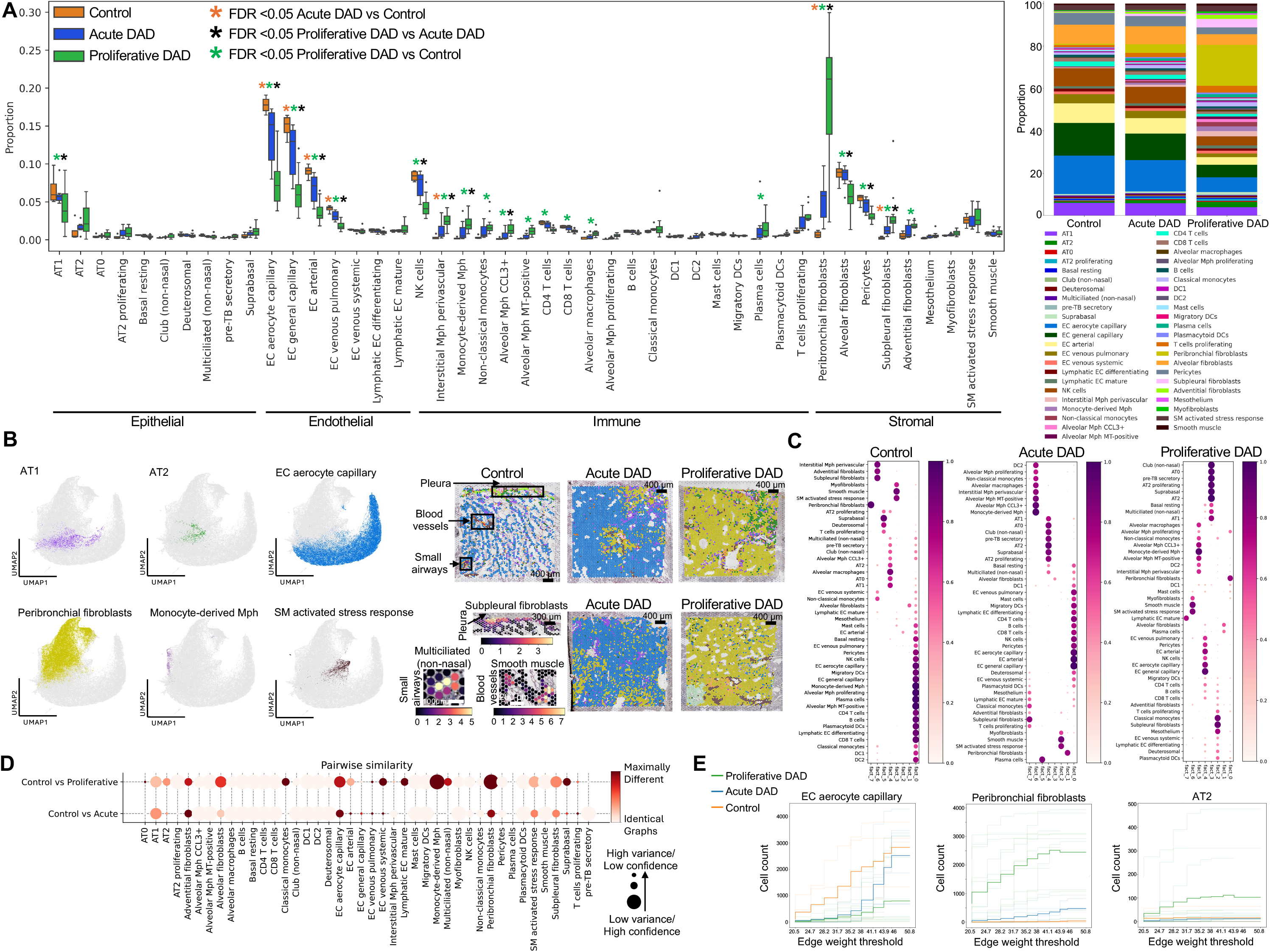
Cell type assignment and cellular compartments. **A)** Bar plot showing the proportion of the 45 cell types identified across disease progression. Significance values indicate credible differences between conditions (FDR < 0.05). **B)** UMAP representation of representative cell types for the different conditions and mapping of cell type deconvolution results on histological images across disease progression. An illustrative example showing cell type density matching expected lung structures is shown. **C)** Cellular compartments identified across disease progression are represented as factors on the x-axis. Cell type loadings are represented by both dot size and color for cell types annotated. **D)** Cell type specific subgraph comparison using the portrait method across condition pairs. Dot size is indicative of the similarity score variance over samples. **E)** Filtration curves for three cell types. A filtration curve is plotted for every sample as well as the mean curve for every condition identified by the thicker and darker line. Large vertical steps towards the left of the plot indicate low density, whereas large vertical steps towards the right of the plot indicate high density.

To assess the spatial distribution of the identified forty-five cell types within neighboring compartments, we applied the cell2location algorithm to the VisiumST data. As expected for the normal lung tissue samples, the characterized cell types mapped within physiological cellular microenvironments such as the great compartment defined by immune and endothelial cells, one rich in epithelial cells (excluding AT2 proliferating, suprabasal and deuterosomal cells that shared a common location), another related to smooth muscle related cells and the fibroblast lineage (where peribronchial fibroblast resided in an isolated compartment) (**Figure 2C**). This compartmentalization underwent an abrupt shift upon DAD progression. The acute DAD stage was characterized by a recruitment of an enriched AT2 proliferating population to the epithelial compartment; the irruption of a spatial cluster of macrophage subtypes and type-2 dendritic cells (DC2s, that promote cytotoxic T-cell responses and helper T-cell differentiation); and the appearance of plasma cells in an isolated population (**Figure 2C**). Most of these cellular redistributions underwent further compartmentalization in the proliferative DAD stage that additionally exhibited the emergence of a unique compartment for lymphatic mature endothelial cells (**Figure 2C**).

To further analyze and characterize tissue architecture differences we also developed an independent methodological approach by applying GraphCompass (Graph Comparison Tools for Differential Analyses in Spatial Systems) (Ali et al., 2024) (**Methods**), a comprehensive set of designed graph analyses methods for “omics” data to quantitatively determine and compare spatial arrangement of distinct cell types among different biological conditions that have been successfully applied to VisiumST data (Ali et al., 2024). In this regard, GraphCompass has been used to evaluate cell-type-specific composition in breast cancer progression, the miocardium following ischemic injury, and for brain development (Ali et al., 2024). Using this approach, we observed the alteration of the spatial organization of cell types from healthy lung to acute and proliferative DAD stages. The cell-type-specific subgraphs across condition pairs, where the size of the dot indicates the pairwise similarity score variances (**Methods**), is shown in **Figure 2D**. Among the characterized distinct cells through COVID-19 progression, we further analyzed peribronchial fibroblasts, endothelial aerocyte capillary cells and AT2 cells by plotting filtration curves for every sample, as well as the mean curve for each lung stage (**Methods**) (**Figure 2E**). These analyses reinforced the findings that these cell types underwent antiparallel shifts in their abundance upon DAD progression: peribronchial fibroblasts and AT2 cells exhibited an overrepresentation whereas endothelial aerocyte capillary cells were depleted, particularly at the proliferative stage (**Figure 2E**).

### Spatial cell-cell interactions in the spectrum of the disease

One of the most exciting applications of spatial transcriptomics is the potential to analyze cell-cell communication (CCC). CCC is a multicellular and complex process involving multiple mechanisms, including intercellular signaling and intracellular signaling as a downstream response associated to the intercellular signaling. Related to the intercellular component, cells interact on diverse levels that include direct contact between ligands and surface receptors, tight junctions, and mechanical forces, and through indirect means, such as the release of soluble factors. For single-cell analyses, the molecular profiles of sender and receiver cell types allow the inference of underlying cell communication events in a tissue using co-occurrence of ligand and receptor (LR) expression among the candidate communicating cells (Browaeys et al., 2020; Efremova et al., 2020) and through gene expression profiles in the receiving cell type related to the extracellular interaction (Arnol et al., 2019; Browaeys et al., 2020). In this regard, most models of intercellular crosstalk depend on the molecular landscape of dissociated cells and, thus, do not pay attention to the location of the studied cells within a tissue. Herein, we used the VisiumST data to identify coordinated cell-cell communication signatures shared across all tissue slides by applying non-negative matrix factorization (NMF) to the estimated local (spot level) ligand-receptor interactions (LRIs), calculated using spatially-weighted Cosine similarity with LIANA+ (Dimitrov et al., 2022; Dimitrov et al., 2023) (**Methods**). Using the elbow selection procedure, we decomposed the local interactions into three Factors (1, 2 and 3) representing three different intercellular communication signatures. The NMF factor scores indicate the strength of each factor in each spot, representing the degree of influence by the associated signature. The averaged factor scores per tissue slide clustered according to lung status are shown in **Figure 3A**. Importantly, Factor 3 distinguished the best between control and COVID-19 associated DAD lung tissues, with high mean scores in proliferative DAD and to a lesser extent in acute DAD (**Figure 3A**). Factor 1 was most prominent in control samples, whereas Factor 2 was more active in a subset of proliferative DAD.

**Figure 3:**
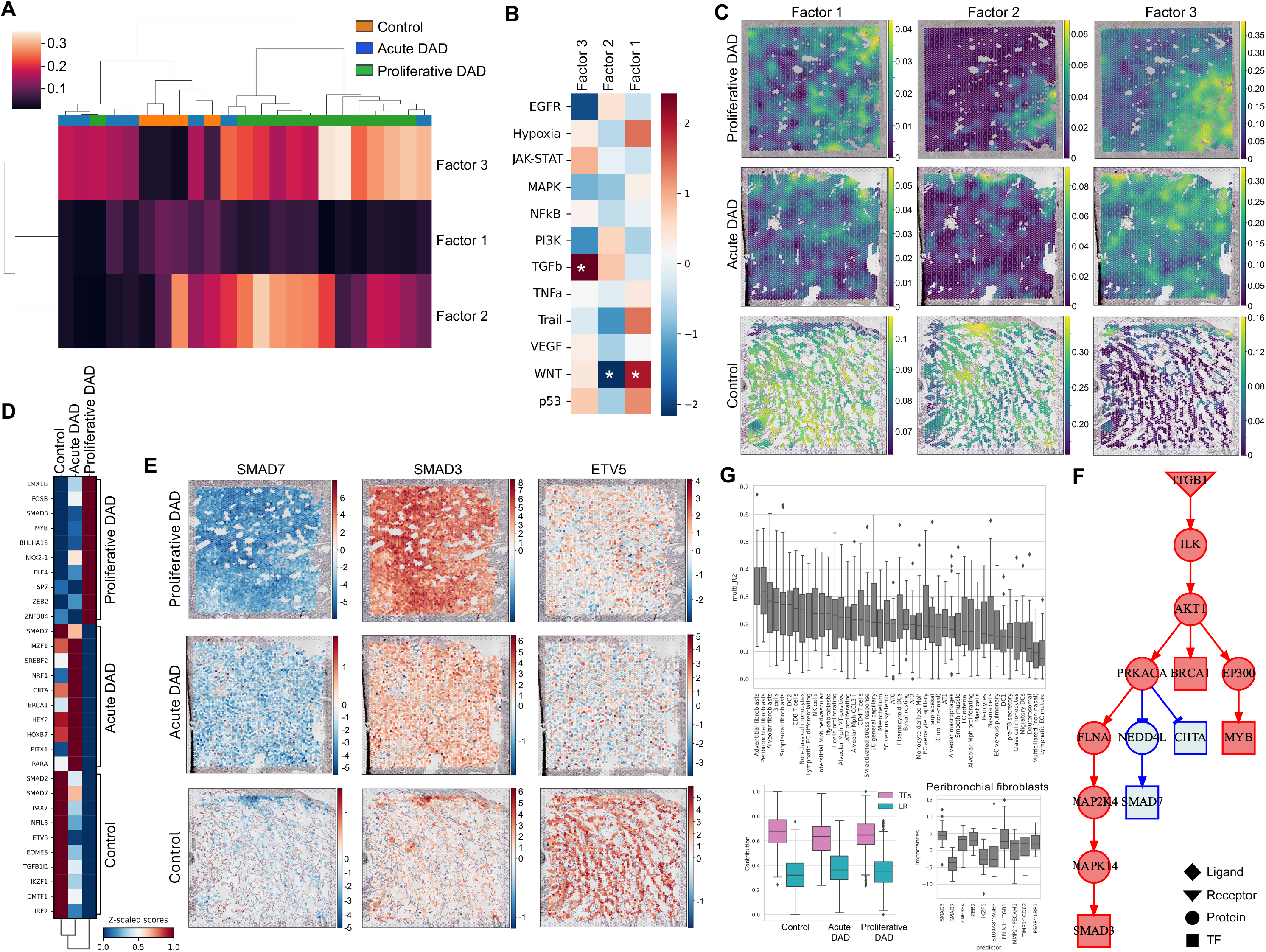
Intercellular and intracellular communication programs in fatal COVID-19. **A)** Heatmap representing average factor scores per lung tissue slide according to ligand-receptor interaction scores. Ward clustering method and Euclidean distance were used to attach samples hierarchical clustering. **B)** Pathway enrichment analysis of ligand-receptor loadings. Statistically significant enrichment scores (p-value < 0.05) are denoted with a star (*). **C)** Factor 1, Factor 2 and Factor 3 scores in selected samples across disease progression. **D)** Heatmap representing transcription factor activity enrichment score. Top 10 transcription factors are shown per condition. Enrichment scores were Z-scaled for comparison purposes. **E)** SMAD3, SMAD7 and ETV5 transcription factor enrichment scores in selected samples across disease progression. **F)** Cell type abundance variance explained by means of R^2^ according to ligand-receptor interactions and transcription factor activity combined predictive performance. Additionally, the contribution of ligand-receptor interactions and transcription factor activity to the predictive performance is shown. Lastly, the top 10 predictors of peribronchial fibroblasts abundance with their corresponding importances as defined by t-values in the predictive model are shown. **G)** Causal intracellular signaling network in peribronchial fibroblasts connecting deregulated intercellular events with downstream transcription factors.

To provide further biological insight, we performed LRIs pathway enrichment analysis on the distinct interaction loadings contributing to the three factors (**Figure 3B**) using multivariate linear regression (Badia-i-Mompel et al., 2022) and pathway annotations from PROGENy (Schubert et al., 2018) (**Methods**). We found that DAD associated Factor 3 was significantly enriched in interactions related to the transforming growth factor beta (TGF-β) pathway (**Figure 3B**), a driver of fibrosis involved in response to tissue injury (Chanda et al., 2019). Conversely, DAD associated Factor 3 was depleted for the EGFR pathway, almost reaching significance (P = 0.055) (**Figure 3B**). Interestingly, the wingless-related integration site (WNT) pathway was enriched in Factor 1 (characteristic of the control samples) but depleted in Factor 2 (**Figure 3B**).

Since Factor 3 was the more optimal discriminator between healthy and COVID-19 associated DAD affected pulmonary tissues, we mainly focused our analyses in this CCC readout. The top four LR loadings defining DAD associated Factor 3 were the interactions TIMP1^CD63, that regulates cell proliferation, survival and migration (Warner et al., 2020), with TIMP1 promoting fibrosis in lung tissues mediated by the TGF-β1/SMAD3 pathway (Duch et al., 2022) being proposed as a potential therapy target (Almuntashiri et al., 2023); APP^CD74, highlighting the role of APP as a lung capillary barrier defense during infection (Vorth, S.B. et al., The FASEB journal 2022) and being associated with failed tissue repair, fibrotic niches and scar-macrophages and natural killers (Ye et al., 2022; Yu et al., 2023); CD99^CD81, regulators of both T-cell and B-cell activity (Pata et al., 2011; Gao et al., 2018); and LUM^ITGB1, with LUM being linked to extracellular matrix (ECM) remodeling and inflammation-associated fibroblasts (Tao et al., 2024) (**Table S1**). Noteworthy additional LRIs involved PSAP (PSAP^LRP1) and members of the S100 protein family S100A8 (S100A8^AGER, S100A8^ITGB2) and S100A9 (S100A9^AGER, S100A9^CD68, S100A9^ITGB2), that have been reported to activate macrophages in COVID-19 (Melms et al., 2021, Rendeiro et al., 2021); the SPARC protein (SPARC^ENG), that promotes microvascular remodeling and act as a downstream effector of TGF-B induced fibrosis (Wong and Sukkar, 2019) and is upregulated in COVID-19-associated fibrosis (Pérez-Mies et al., 2022, Melms et al., 2021) and idiopathic pulmonary fibrosis patients (Conforti et al., 2020;); and vimentin (VIM^CD44), an important attachment factor for SARS-CoV-2 entry into endothelial cells that contribute to COVID-19 vascular complications (Amraei et al., 2022). Beyond the mentioned ligand proteins, it is also relevant to mention that the three most frequent receptors involved in the LRIs (**Table S1**) were CD44, involved in T-cell abundance and fostering of the cytokine storm linked to poor prognosis of COVID-19 patients (Zick, 2022); ITGB1, that associates with the angiotensin-converting enzyme 2 (ACE2) to mediate SARS-CoV-2 entry (Zhang et al., 2022) and regulates ECM remodeling; and LRP1, that is involved in the overproduction of cytokines and chemokines (Zick, 2022) and enriched in those COVID-19 patients that died (Razaghi et al., 2022) or underwent long haul COVID-19 disease (Gu et al., 2023). The NMF factor scores indicating the strength of each factor in each spot are depicted for illustrative examples of VisiumST slide images throughout COVID-19 induced DAD progression and shown in **Figure 3C**.

Regarding intracellular signaling analysis, we studied transcription factor activities including their downstream transcriptional targets that shift in the progression from normal lung to the COVID-19 associated acute phase and, at the end, to the proliferative DAD stage. We estimated transcription factor (TF) activity in each VisiumST spot based on multivariate linear regression using decoupleR (Badia-i-Mompel et al., 2022) and CollecTRI (Müller-Dott et al., 2023) network containing a curated collection of TFs and their targets (**Methods**). **Table S2** shows the TFs activity in each type of lung tissue and **Figure 3D** displays the top 10 Z-scaled TF enrichment scores in each condition that discriminate between control lungs and the pulmonary samples of COVID-19 associated acute and proliferative DAD. One important observation is that these data highlight the critical role of TGF-β pathway in DAD progression, as did also the intercellular LRI analysis (**Figure 3B**). In this regard, it is noteworthy to mention the opposite activity landscape of SMAD protein family members. The COVID-19 associated proliferative DAD stage is characterized by upregulation of SMAD3 activity, a key mediator of TGF-β signaling to promote ECM production, tissue repair, fibrosis and scar formation after injury (Finnson et al., 2010). Conversely, SMAD7 activity that exerts antagonizing roles to TGF- β/SMAD3 profibrotic pathway, is downregulated across DAD progression. Illustrative VisiumST slides depicting the described TF activity patterns of SMAD TFs in COVID-19 associated DAD progression are shown in **Figure 3E**. The altered activity of the TGF-β signaling pathway in the natural history of the disease was also further strengthened by the involvement of two additional components, TGFB1|1 [a marker of contractile smooth muscle cells (Wang)] and ZEB2 [epithelial to mesenchima transition and fibrogenesis (Teraishi et al., 2017)], that were upregulated in control and proliferative DAD lungs, respectively (**Figure S2**). Interestingly for the last gene, ZEB2 DNA methylation status has been linked to another severe consequence of SARS-CoV-2 infection, the multisystem inflammatory syndrome in children (MIS-C) (Davalos et al., 2022). In addition to TGF-β signaling, another two cellular networks were targeted by aberrant TF activity: lung epithelial cell differentiation and NK cells functionality. In the first case, downregulation of the ATII cell identity regulator ETV5 occurred upon COVID-19 induced DAD progression as shown in **Figure 3E**, suggesting initiation of epithelial regeneration by ATII cells (Melms et al., 2021; Zheng); whereas NKX2-1 [regulator of alveolar epithelial progenitors (Toth et al., 2023)], MYB [involved in airway epithelial cell differentiation (Pan et al., 2014)] and BHLHA15 [linked to acinar cell function (Ref)] were upregulated in proliferative DADs (**Figure S2**). Remarkedly, SREBF2, related to surfactant production in ATII cells, and CIITA, which drives MHCII expression and induces cell resistance to SARS-CoV-2 (Bruchez et al., 2020), was enriched in acute DAD lungs (**Figure S2**). For NK cells, where we already observed a quantitative decrease in COVID-19 progression (**Figure 2A**), multiple TFs essential for their proper development were also downregulated in the DAD lungs, such as IRF2 (Persyn et al., 2022), IKZF1 (Goh et al., 2024), NFIL3 (Gascoyne et al., 2009) and EOMES (Zhang et al., 2021) (**Figure S2**), supporting that the proposed NK cell dysfunction in COVID-19 (Bi, 2022) could be associated to fatal forms of the disease.

### Spatial relationships of ligand-receptor interactions and transcription factor activities with cell type abundance

Using VisiumST data, we have above provided a comprehensive assessment of the targeted cell types, the intercellular and the intracellular signaling that characterizes the consecutive steps from a morphologically normal lung to the tissue affected in the COVID-19 induced acute DAD phase to the late proliferative stage. To further capture how local LRIs, TFs and the distinct cell type abundances relate in the lung spatial context, we leveraged an explainable multi-view modelling approach to decipher the global spatial relationships between these three components (Tanevski J. et al., 2022). In this regard, we jointly modelled, in a spatially informed manner, cell type abundances in each spatial spot using the top 25 local ligand-receptor loadings from Factor 3 (**Table S1**) and the activity of the top 10 most enriched TFs (**Table S2**) per condition. We observed that across control and DAD lung tissue slides, both LRIs and TFs activity jointly contributed to explain the variance of cell type abundance (mean multi-view R^2^ = 0.21) (**Figure 3F**). The relative contribution of each spatial “view” to the joint predictive performance was higher for TF activities (median contribution > 60%) compared to LRIs (median contribution < 40%) across disease progression (**Figure 3F**). A high degree of variability in variances explained across tissue slides and cell types was also observed. In this regard, the abundance of fibroblasts was best explained by these components [including activity of TFs SMAD3 and SMAD7 and TIMP1^CD63 LRI among top 10 predictors of peribronchial fibroblasts (**Figure 3F**)], highlighting their contribution to DAD. Additionally, important differences were also found when explaining cellular composition across control lungs and COVID-19 induced DAD progression, particularly for peribronchial fibroblasts, endothelial aerocyte capillary cells, endothelial general capillary cells, alveolar macrophages subtypes, monocyte-derived macrophages, non-classical monocytes and NK cells (**Figure S3**). These last results further highlight the critical role of these cell types in COVID-19 induced DAD lung remodeling reflected by the observed temporal differences.

Additionally, we used the predictor importances (coefficients’ t-values) from this predictive linear model to infer intracellular signaling networks linking both LRIs and TF activity patterns. To accomplish this aim, we applied LIANA+ (Dimitrov et al., 2023) (**Methods**), to infer a putative causal network linking LRIs to TFs using a network optimization approach. To better understand the pathological lung tissue repair response in DAD, we inferred intracellular signaling pathways focusing on peribronchial fibroblasts considering their significant enrichment in proliferative DAD lungs (**Figure 3G**) and in idiopathic pulmonary fibrosis patients (Madissoon et al., 2023). Our analyses suggested that initial activation of the ITGB1 receptor unleashed a downstream signaling pathway that caused an upregulation of the transcription factors SMAD3, MYB and BRCA1 and a downregulation of SMAD7 and CIITA (**Figure 3G**). Importantly, CIITA represses collagen expression by lung fibroblasts after injury (Xu et al., 2011). Interestingly, and beyond a better basic understanding of the pathophysiology of the disease, this pathway analysis shows that integrin ITGB1 induces an activation of the kinases ILK, AKT1, MAP2K4 and MAPK14 that have been previously linked to the disorder (Bouhaddou et al., 2020; Xia et al., 2020) and could be now considered as even more amenable candidates for new targeted therapies.

The characterized global spatial relationships are calculated at the level of the whole VisumST slide. To identify local spatial dependencies that might occur only in a sub-region of the studied lung tissues and to pinpoint their precise location, we leveraged spatially-informed local bivariate similarity metrics, that included spatially-weighted Cosine similarity and global Morańs R (**Figure 4A**), to identify pairs of LRIs that are spatially clustered together or apart (**Methods**). The LRI TIMP1^CD63 showed the highest spatial co-clustering pattern by both Cosine similarity and globan Moran’s R metrics (**Figure 4A**). The pattern of the co-clustering was particularly evident in proliferative DAD lungs within DAD associated Factor 3 boundaries (**Figure 4B and Figure S4**). Computed permutation-based p-values to assess the significance of the local interactions demonstrated an agreement with the high Cosine similarity regions (**Figure 4B**). To further categorize TIMP1^CD63 spatial relationship, we identified that for the majority of local category areas, both ligand and receptor were highly expressed and only in a few sub regions one interaction member was highly and the other lowly expressed (**Figure 4B**). Interestingly, the amyloid-beta precursor protein (APP) participated in multiple LRIs, including APP^CD74 with the second highest Cosine similarity, albeit a weak global Morańs R (**Figure 4A**). An additional APP LRI, APP^AGER, showed a clear distinct and diminishing spatial co-clustering pattern over disease progression (**Figure S4** **and** **Figure S5**), suggesting a meaningful biological relationship since AGER is an AT1 marker described in the literature (Madissoon et al., 2023) and also reported in our study (**Figure S1**). Intriguingly, beta-amyloid produced by the infection-mediated lung injury can reach through general circulation other organs originating further defects, including neurocognitive dysfunction (Balczon et al., 2024). Strikingly, cognitive impairment in the post-acute phases of COVID-19 is not an uncommon observation (Wang et al., 2024) and SARS-CoV-2 infection is considered a risk factor for Alzheimeŕs disease (Bonhenry et al., 2024). Finally, multiple LRIs with proteins involved in the activation of Mphs in COVID-19, such as S100A9, showed increased spatial co-clustering through disease progression (**Figure S4** **and** **Figure S6**). Interestingly, S100A9^CD68, the best LRI predictor of all Mph subtypes, non-classical monocytes and T cells proliferating cell type abundances (**Figure S6** **and** **Figure S7**), yielded one of the highest median global Morańs R scores (**Figure 4A**), suggesting an important role contributing to aberrant myeloid activation and dysregulated immune response (Melms et al., 2021; Merad et al., 2022). The mapping of the described LRIs on the VisiumST images in illustrative control and COVID-19 associated acute and proliferative DAD cases are shown in **Figure 4B**, **Figures S5** and **S6**.

**Figure 4:**
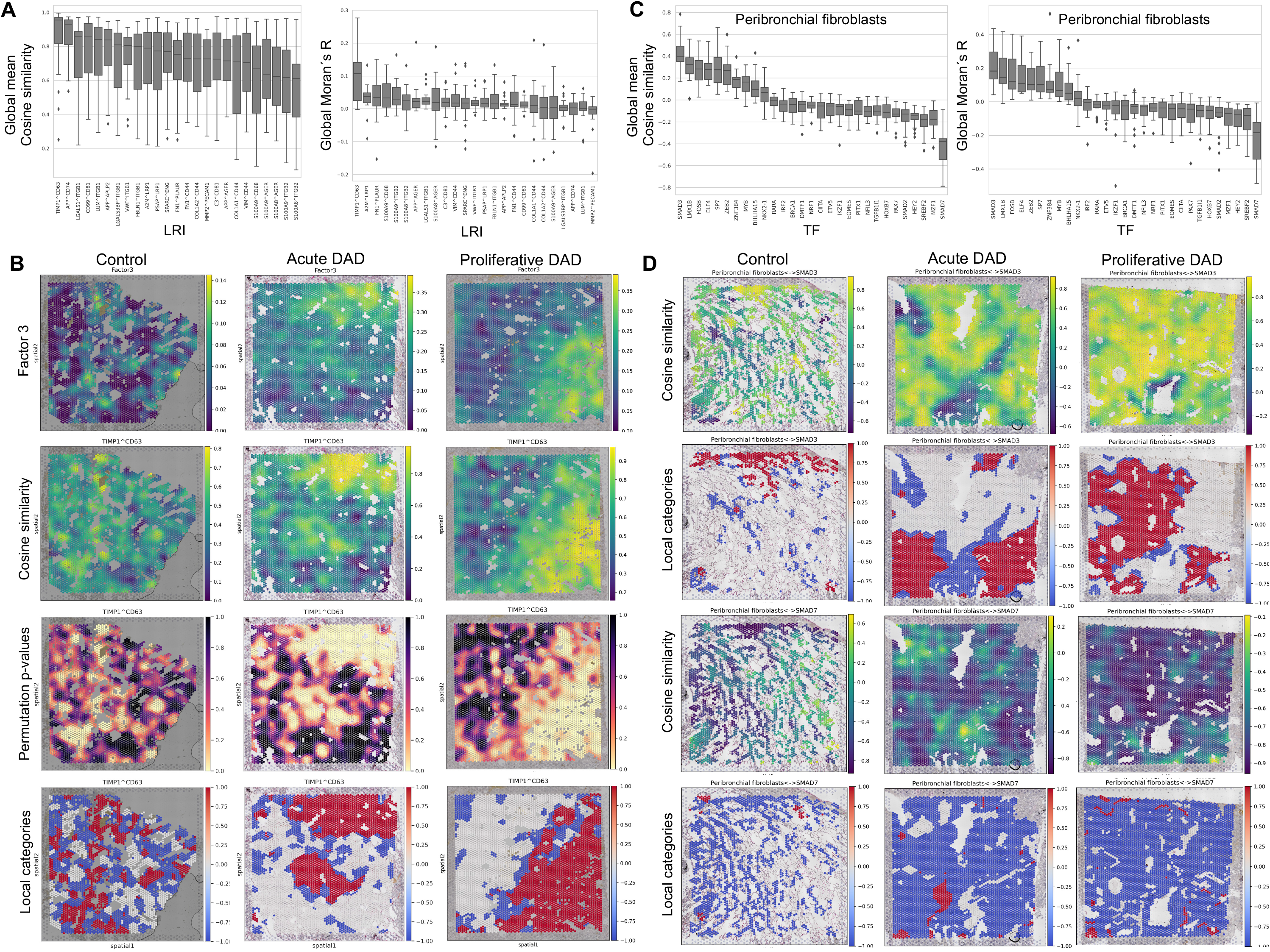
Spatial local dependencies in ligand-receptor interactions and between cell types and transcription factor activity. **A)** Local bivariate similarity metrics score, including spatially-weighted cosine similarity and bivariate Morańs R for the top 25 ligand-receptor loadings defining Factor 3 in all studied samples. **B)** Mapping of Factor 3 scores, cosine similarity, permutation based p-values and local categories for TIMP1^CD63 interaction in selected samples across disease progression. **C)** Local bivariate similarity metrics score, including spatially-weighted cosine similarity and bivariate Morańs R for the top 10 transcription factor activity and peribronchial fibroblasts abundance in all studied samples. **D)** Mapping of SMAD3 and SMAD7 local interactions with peribronchial fibroblasts according to cosine similarity and local categories in selected samples across disease progression.

In a similar manner that we analyzed the local spatial dependencies for LRIs, we next leveraged the spatially-informed local bivariate similarity metrics to investigate associations between cell types and TFs activity considering their relevance for tissue function, using also spatially-weighted Cosine similarity and global Morańs R (**Figure 4C**) (**Methods**). Peribronchial fibroblasts (PBFs), the most abundant cell type in COVID-19 associated DAD proliferative lungs (**Figure 2A**), were most spatially associated and co-clustered with the pro-fibrotic TF SMAD3 activity locations (**Figure 4C**, **4D**), whereas PBFs and the anti-fibrotic TF SMAD7 activity locations were spatially clustered apart (**Figure 4C**, **4D**). These results fit the previously characterized mutually exclusive location of SMAD3 and SMAD7 activities through DAD progression (**Figure 3E**) and the top LRI loading, TIMP1^CD63, characterizing DAD associated Factor 3 in COVID-19 induced DAD tissues promoting lung fibrosis through the TGF-β1/SMAD3 pathway (**Figure 4B**). All these results highlight the central role of TGF-β pathway activation in driving pathological ECM remodeling and repair linked to aberrant activation of PBFs that leads to scar formation and a grossly disrupted lung tissue architecture in the COVID-19 associated DAD proliferative cases.

To provide a second example beyond PBFs of local spatial relationships between TFs activity and the identified cell types, it is worth highlighting the myeloid lineage. We observed that the activity of the TF MYB was the best predictor of the abundance of myeloid cell types (**Figure S7**). The spatial co-expression of MYB with the myeloid and T cell proliferating cell fates increases as disease progresses (**Figure S8**). MYB plays an essential role in many hematopoietic pathways (Ref) and, most importantly, the E2F/MYB regulatory programs from myeloid cell populations have been recently described as hyperactivated in COVID-19 patients with poor disease outcomes (Lam et al., 2023), that in a similar manner we have herein observed with a spatial perspective. The interrogation of the immune landscape by assessing local relationships between TFs activity and cell types also unveiled that NK cells, another population critically depleted in our COVID-19 associated DAD cases (**Figure 2A**), were most influenced by IKZF1 activity showing the highest co-clustering pattern (**Figure S8**). IKZF1 is essential for proper NK cell development (Goh et al., 2023). Here we report a loss of the co-clustering pattern of IKFZ1 and NKs across COVID-19 associated DAD progression (**Figure S8**). Results that strengthen the suggested central role of NK cell dysfunction in the development of fatal COVID-19 (Krämer et al., 2021).

### Cell-cell communication as a function of niche composition

The potential cell-cell communication events that could occur in lung tissues across the different conditions was assessed not only accounting for LRIs and TFs activity, as explained above, but using a graph neural network method (NCEM) (Fischer et al., 2023) that estimates the effect of the inferred spot composition on gene expression variation within cell types across spots to discover intercellular dependencies (‘**Methods’**).

To discriminate different intercellular dependencies between control and fatal COVID-19 lung sections, we identified multiple cell type couplings across the most abundant and variable cell types over disease progression (**Methods)** observing a profound reconfiguration of intercellular communication (**Figure 5A**). We observed a dependency of NK cells in control lung tissues on various cell types, including CD8 T cells, EC aerocyte capillary cells, alveolar Mph CCL3+ and AT1 cells. However, these dependencies were lost in acute and proliferative COVID-19 associated DAD lung tissues (**Figure 5A**). Most importantly, we observed that in DAD samples the population of CD4 T cells beared multiple dependencies on various cell types such as non-classical monocytes, SM activated stress response cells, plasma cell and AT1 cells, becoming a prominent receiver node of communication, particularly in acute DAD and to a lesser extent in proliferative DAD lungs. Interestingly, AT1 cells exhibited limited intercellular dependencies in control lung samples, that increased in the acute DAD phase, and were lost in the proliferative DAD stage. Lastly, CD4 T cells established a dependency on AT2 cells in the proliferative DAD phase. Additionally, we performed a receiver effect analysis highlighting gene-wise effects of all senders on once receiver cell type to contextualize gene expression differences in some of the couplings (**Figure 5B**). Furthermore, since CD4 T cells showed important dependencies on multiple cell types in DAD lungs, we performed a sender similarity analysis to characterize the profile of these intercellular dependencies across DAD progression. We observed that in acute DAD lungs, the sender profile mostly conserved lineage cell type identity but was lost in proliferative DAD lungs (**Figure 5B**).

**Figure 5:**
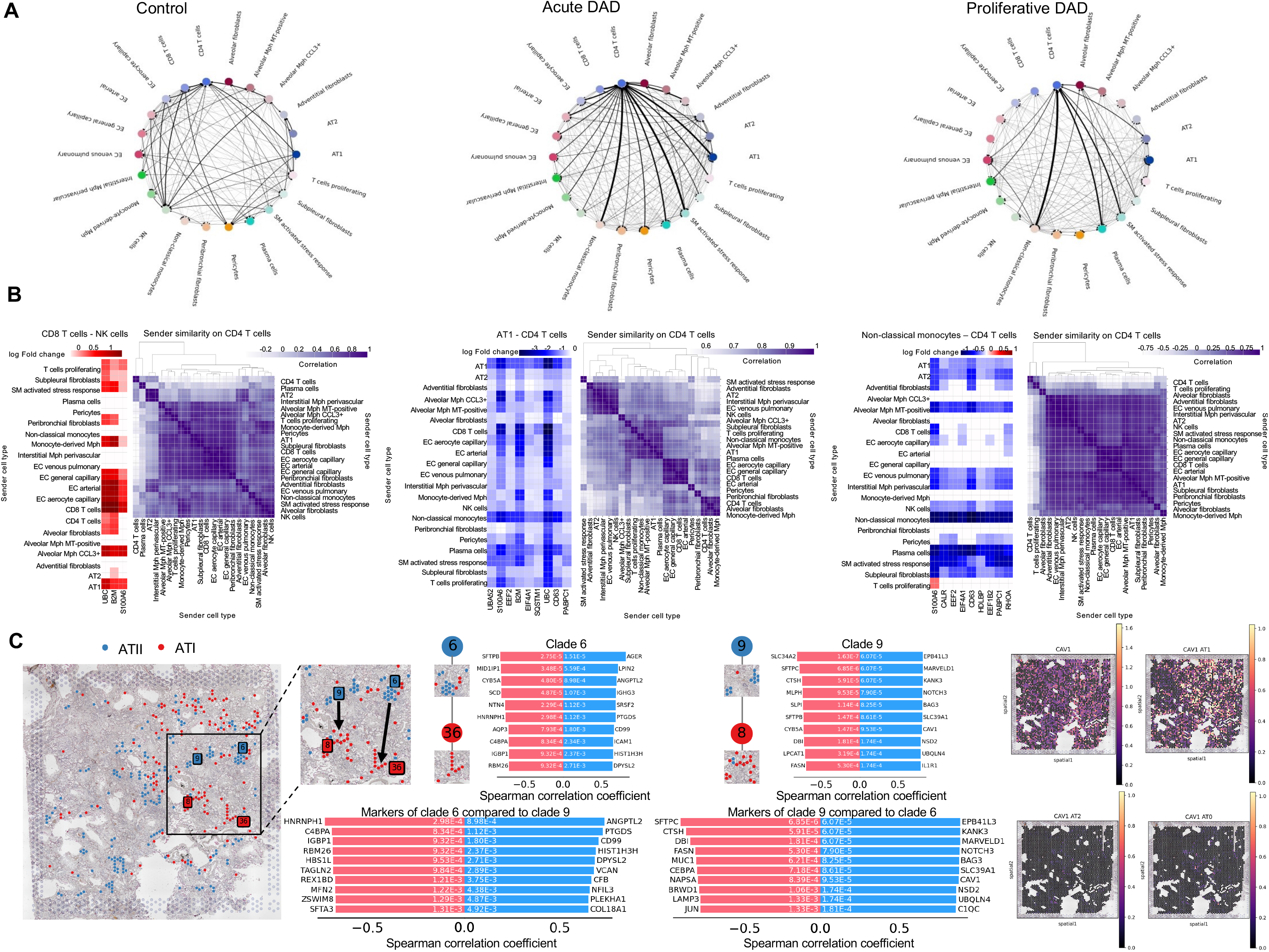
Intercellular dependencies as a function of niche composition and spatio-temporal trajectories. **A)** Type coupling analysis with edge proportional to strength of directional dependencies by means of fold changes of differentially expressed genes for each pair of sender and receiver cell types. Only edges with at least 500 genes are shown. Results for intercellular dependencies across disease progression are shown. **B)** Sender effect analysis of the CD8 T cells – NK cells axis in control samples, AT1-CD4 T cells axis in acute DAD lungs, and non-classical monocytes and CD4 T cells in proliferative DAD lungs. Shown is the estimated fold change that the sender cell type on the y-axis induces in the gen on the x-axis in receiving cells. Additionally, a sender similarity analysis based on a correlation of the coefficient vectors of each sender type with respect to CD4 T cell receivers across disease progression is shown. **C)** Spatio- temporal trajectory of AT2 to AT1 cell type differentiation in a proliferative DAD lung tissue identified two clades of transdifferentiating cells. Transition genes positively (blue) or negatively (red) correlated with the predicted trajectory and extracted by Spearman correlation test with adjusted p-value <0.05 and correlation coefficient > 0.4 or < -0.4 are shown for each transitioning clade. A comparison between clade markers is additionally shown. Lastly, total and cell type specific gene expression of CAV1 is shown for AT1, AT2 and AT0 cells.

### Spatio-temporal trajectories: AT2-AT1 epithelial regeneration

AT2 cells play an important role as AT1 progenitors during lung injury, proliferating and contributing to alveolar repair and regeneration. By contrast, AT1 cells are fragile, susceptible to damage and unable to proliferate. Therefore, characterizing AT2-AT1 differentiation is important to provide novel insights into cellular processes and tissue repair mechanisms in severe COVID-19. To decipher the dynamic relationships across tissue space and time between transcriptional states of AT2 and AT1 cells, we leveraged a spatial graph-based method named pseudo-time-space (PSTS) implemented in the stLearn software (Pham et al., 2023) ‘**Methods**’). By combining spatial and imaging information, representing cell and tissue morphology, with gene expression data, we used the PSTS algorithm to map the spatial changes in AT2 and AT1 cell states, modelling and reconstructing their spatio-temporal trajectories. In this regard, we defined a spatial trajectory for AT2 cells transitioning into AT1 cells across two clusters (clade 6 and clade 9) in a proliferative DAD lung (**Figure 5C**). The top 10 most upregulated and downregulated genes defining AT2 to AT1 transition in each clade are shown (**Figure 5C**). Enrichment analysis (’**Methods**’) revealed that the top 10 upregulated genes in clade 6 were enriched (FDR q-value = 0.001) in AT1 cell identity markers including ICAM1, DPYSL2, ANGPTL2 and AGER (Travaglini et al., 2020). Likewise, the top 10 downregulated genes in clade 6 were enriched (FDR q-value < 0.001) in AT2 cell identity markers including SFTPB, SCD and CYB5A (Travaglini et al., 2020). Furthermore, the top 10 downregulated genes in clade 9 were also enriched (FDR q-value < 0.001) in AT2 cell identity markers including SFTPB, SFTPC, SLC34A2, FASN, CTSH, DBI, MLPH, LPCAT1 and CYB5A (Travaglini et al., 2020). These results further validate the inferred spatio-temporal trajectories in AT2 and AT1 cell states, better characterizing the alveolar epithelial regeneration process after lung injury. Interestingly, when comparing clade 6 and clade 9 (**Figure 5C**), we found that CAV1, a late AT1 maturation marker (Melms et al., 2021) was upregulated in clade 9, suggesting a complete transition of AT2 to AT1 cells. This is further supported by the cell type specific gene expression of CAV1 in spatial coordinates among different subtypes of pneumocytes, where CAV1 is expressed only by AT1 cells in distinct locations but not by co-located AT2 and derived AT0 cells during alveolar repair, driving most of CAV1 total expression across all cell types (**Figure 5C**). Altogether, these results reinforces the notion that, in the lung of patients with severe COVID-19, AT2 cells aim to repopulate AT1 cells upon the activation of alveolar epithelial regeneration programs.

## Discussion

COVID-19 exhibits a great range of clinical behaviors, with most of SARS-CoV-2 infected individuals displaying only a few symptoms or being completely asymptomatic (Coronaviridae Study Group, 2020; Wu et al., 2020). However, the natural history of the disease could also end with the death of the patient, particularly when severe respiratory failure occurs linked to acute respiratory distress syndrome and interstitial pneumonia. The causes behind the wide spectrum of clinical manifestations of COVID-19 are mostly unknown, although life-threatening COVID-19-associated respiratory failure occurs more frequently in aged males with concomitant medical conditions, such as obesity, diabetes, hypertension and cardiovascular disease (Zhou et al., 2020; Li et al., 2020). Several genomic and epigenomic biomarkers have also been associated with COVID-19 life-threatening pneumonia (Covid-19 GWAS Group Severe et al., 2020; Pairo-Castineira et al., 2020; Zhang et al., 2020; Bastard et al., 2020; Castro de Moura et al., 2021). Focusing on one of the main target tissues of the disorder, the lung, transcription profiles in extracts or laboratory-suspended cells from autopsy specimens of pulmonary samples have also shown at both bulk transcriptome level (Blanco-Melo et al., 2020; Pinto et al., 2020; D’Agnillo et al., 2021) and high-resolution single-cell transcriptional magnification (Mels et al., 2021; Delorey et al., 2021; Sikkema et al., 2023) some of the expression profiles tied to fatal COVID-19. However, this wealth of information is derived from disaggregated lung tissues where the architecture of the organ and its potential shift in the disease is not preserved. This can be extremely important to understand the pathophisiology of COVID-19 because RNA expression patterns do not only depend on internal gene regulation, but also the influence of neighboring cells and, overall, on the surrounding tissue microenvironment. This issue has been scarcely addressed in COVID-19 affected lung tissues (Desai et al., 2020; Margaroli et al., 2021; Park et al., 2022; Mothes et al., 2023). To solve these issues, we have herein applied in the lung of fatal COVID-19 patients the spatial transcriptomic technology that delivers the transcriptional pattern of cells by RNA-sequencing preserving the organization of the tissue in the organ. Importantly, spatial transcriptomics can be combined with microscopic imaging and immunohistochemistry, as also herein described, to improve our understanding of where even further in the spatial context these expression changes are taking place. The picture that emerges is one defined by profound shifts in specific cellular populations of the epithelium, endothelium, fibroblasts and immune cells with distorted intercellular communications that finally disrupt important gene networks leading to fatal outcomes.

The disbalance between cell types exhibited the most remarkable change for peribronchial fibroblasts that experimented an extraordinary increase in the proliferative stage of the disease. Other subclasses of fibroblasts also underwent upregulation among COVID-19 progression, except for alveolar fibroblasts that decreased in a similar fashion of other cell types populating the functional respiratory alveolus such as epithelial AT1 cells and aerocyte capillary endothelial cells. Interestingly, the proliferative DAD phases showed an increase in various subtypes of alveolar macrophages. Furthermore, the myeloid lineage underwent an overall increase, including interstitial and monocyte-derived macrophages and non-classical monocytes, whereas the lymphoid lineage decreased across DAD progression, mostly driven by the loss of NK cells in proliferative DAD lungs. These results match the reported aberrant activation of myeloid cells and impaired T cell and NK cell responses in fatal COVID-19 (Melms et al., 2021; Krämer et al., 2021). In this regard, we highlight the important role of NK cells in severe COVID-19, characterizing NK cells dysfunction with marked downregulation of essential TFs activity for their functional development and maturation, including IKZF1 (Goh et al., 2024), IRF2 (Persyn et al., 2022), EOMES (Zhang et al., 2021) and NFIL3 (Gascoyne et al., 2009). Herein, we report a progressive and persistent dysfunction of NK cells throughout DAD spectrum, further implicating NK cell functional impairment in promoting lung fibrosis in fatal COVID-19 (Krämer et al., 2021). Besides NK cells dysfunction, a dysregulated immunological repair response to SARS-Cov-2 infection has been proposed as a major contributor to disease progression (Merad et al., Science 2022). Hence, we have described the LRIs and TFs activity related to essential elements of the immune system, identifying S100A9^CD68 LRI and MYB TF activity as major determinants of myeloid cell types abundance, showing increased spatial co-expression patterns over DAD progression, highlighting their important role in the aberrant activation of macrophages in severe COVID-19 (Rendeiro et al., 2021; Lam et al., 2023).

Moreover, we have characterized the molecular drivers of pathological responses to lung injury leading to massive fibrosis and grossly disrupted tissue architecture. We report the key role of TGF-β pathway in DAD progression, identifying an upregulation of the profibrotic SMAD3 activity coupled with downregulation of antagonizing SMAD7 activity. Importantly, we have identified TIMP1^CD63 LRI as a major contributor to DAD, emphasizing TIMP1 role as a key regulator of ECM homeostasis and downstream effector of TGF-β pathway activation, being identified as a candidate therapy target for pulmonary fibrosis (Almuntashiri et al., 2023). Furthermore, when connecting deregulated intercellular communication events to downstream intracellular signaling pathways, we inferred an intracellular signaling network in PBFs suggesting that the phenotypic changes and the different targeting of the SMAD TFs involved the activation of integrin ITGB1 receptor and their associated downstream kinases AKT1, MAP2K4 and MAPK14, representing additional potential targets for COVID-19 therapies (Bouhaddou et al., 2020; Xia, Qui-Dong, et al., Cell Proliferation 2021).

Interestingly, when analyzing intercellular dependencies as a function of niche composition, we described a dependency of NK cells on various cell types in control lungs that was lost on DAD lungs, whereas a dependency of CD4 T cells on multiple cell types including CD8 T cells in control lungs shifted towards other lymphoid and myeloid immune cells and stromal cells in DAD, such as non-classical monocytes, plasma cells and smooth muscle activated stress response cell types, including epithelial AT1 cells in acute DAD and AT2 cells in proliferative DAD, further reflecting the disrupted tissue architecture across the DAD spectrum.

Importantly, our spatio-temporal trajectories analysis helps to characterize the alveolar epithelial regeneration process, highlighting the important role of AT2 cells as AT1 progenitors and identifying markers of AT2 to AT1 differentiation. This process is impaired in severe COVID-19 induced DAD cases, contributing to fatal outcomes in these patients. Following these last observations, it is tempting to speculate that the occurrence of long COVID-19 could relate to the partial or complete disruption of intercellular communication events and differentiation trajectories that cannot completely restore the functional alveolar gas exchange capacity and/or prevent the persistence of fibrotic scars.

Our findings could also provide and foster the research of small drugs and antibodies targeting some of the cell types, pathways and intercellular communication that characterize the spatial aftermath of severe COVID-19. One example could be related to the combating of the fibrosis associated with DAD progression. Pirfenidone and nintedanib are two antifibrotic drugs approved for the treatment of idiopathic pulmonary fibrosis (IPF) (Sing and Wairkar, 2024; Perez-Favila et al., 2024; Li et al., 2024). These agents can also be repurposed to avoid severe COVID-19 associated fibrosis since Pirfenidone inhibits TGF-β (Sansores, et al., 2023), the main profibotic pathway underpinned in our study, and nintedanib blocks several tyrosine and serine/threonine kinases (Landi et al., 2020; Umemura et al., 2021) among them the MAPKs and AKT1 identified in our intracellular pathways analyses of peribronchial fibroblasts. Furthermore, peribronchial fibroblasts are enriched in idiophatic pulmonary fibrosis patients (Madissoon et al., 2023), suggesting common mechanisms underpinning both idiopathic pulmonary fibrosis and DAD diseases. Moreover, we suggest MAPK14 kinase activation role within the intracellular signaling pathway of peribronchial fibroblasts leading to SMAD3 activation. Importantly, MAPK14 inhibitors such as ralimetinib, clinically tested in patients with advanced cancers (Patnaik et al., 2016), and ARRY-797, clinically tested in patients with dilated cardiomyopathy (Judge et al., 2022), have shown SARS-Cov-2 antiviral activity (Bouhaddou et al., 2020), being candidates for drug repurposing in severe COVID-19. Interestingly, several inhibitors of the integrin αvβ6, such as GSK3335103 and BG00011 are also at different levels of preclinical studies or even in clinical trials to treat idiopathic pulmonary fibrosis. Following this lead, our finding that integrin ITGB1 activation unleashes a profibrotic signaling in peribronchial fibroblasts in fatal COVID-19 cases suggests that this receptor can be also another amenable target to avoid the complications of the disorder. A similar case can be drawn for the therapeutic counteract of the emerging and hyperactivated population of inflammatory myeloid cells that we have observed in our spatial transcriptomic analysis. In this regard, macrophages and monocytes engage the NOD-like receptor family pyrin domain containing 3 (NLRP3) inflammasome during SARS-CoV-2 infection and established COVID-19 (Lécuyer et al., 2023). Thus, in addition to the use of general anti-inflammatory interventions, our results further supports that the application of NLRP3 inhibitors, such as NT0793/NT0249 or MCC950 (Diarimalala et al., 2023), can be another useful strategy to restrict the activity of these belligerent myeloid cells and treat severe COVID-19 cases. Importantly, because our integrative spatial transcriptomics analysis of TFs activity within the identified cell types yielded MYB as a major determinant factor for macrophages in severe COVID-19 associated DAD lungs, it could be proposed as a candidate drug target (Uttarkar et al., 2017; Samy et al., 2022). Similarly, we report S100A9^CD68 LRI as a major determinant of myeloid cell types. Interestingly, S100 protein family members S100A8 and S100A9 have been proposed as potential biomarkers for COVID-19 severity, modulators of the cytokine storm and have been investigated as potential targets of small molecules such as Paquinimod to control aberrant myeloid activation in severe COVID-19 (Mellett et al., 2021; Guo et al., 2021).

Related to treatment, we can also briefly mention a provocative thought. Since multiple LRIs including our second ranked LRI according to Cosine similarity co-clustering metric involved the Amyloid-Beta Precursor Protein (APP^CD74), and it has been proposed that common cognition defects in post-COVID-19 (Wang et al., 2024) could be due to beta-amyloid produced by the lung lesions and liberated to the blood (Balczon et al., 2024), including that SARS-CoV- 2 infection is considered a risk factor for Alzheimeŕs disease (Bonhenry et al 2024), we suggest that maybe we can target both processes: the lung injury and the associated cognitive impairment. Following this train of thought, the lysophosphatidic acid receptors (LPARs) from the G protein-coupled receptor family contribute to both Alzheimer’s disease (AD) and bind to the viral SPIKE protein being implicated in COVID-19 inflammation, and LPAR inhibitors are started to be explored for this potential double effect (Malar et al., 2024).

Our study is unique because we provide a spatially informed characterization of the cellular and molecular hallmarks of lung tissue architecture in fatal COVID-19. This detailed spatial transcriptomics study that highlights *in situ* the disease-associated changes in the composition of cellular subsets, their spatial dependencies and disrupted intercellular communication programs also constitutes a proof-of-principle of the potential translational use of the emerging spatial technologies. This transition will require careful benchmark comparison studies among the competing spatial transcriptomic platforms, harmonization of data processing pipelines and design of user-friendly databases where the data can be deposited and interrogated, automatization of sample processing and data analysis workflows leading to a shorter timeframe to deliver the results together with the ongoing reduction of sequencing costs; and scalable computational methods to exploit spatial transcriptomics data. Related to this last point, spatial transcriptomics can constitute one of the entrance points for the application of artificial intelligence in pathology and modern medicine. In this regard, our investigation of the altered cellular and molecular architecture of the lung in fatal COVID-19 could serve as an excellent example of the versatility of spatial transcriptomics to fulfill the promise of how the new genomic technologies could improve our understanding and the personalized management of many human diseases.

## Contributors

C. A. G.-P. provided the bioinformatic analyses of the spatial transcriptomics data. E. M., V. D. and G. F. reviewed the clinical data. X. S. S., E. E., B. P. M., T. C. C., J. Pal. and E. M. reviewed the postmortem lung samples. E. M. performed the immunostainings. M. E. designed the study and wrote the manuscript with contributions from all authors. The study was approved by the institutional ethical review boards of Ramón y Cajal University Hospital (Necropsias_Covid19; 355_20) and the Lund Hospital (ref 2020- 02369).

## Data sharing

## Acknowledgements

We thank CERCA Programme / Generalitat de Catalunya for institutional support. The Secretariat for Universities and Research of the Ministry of Business and Knowledge of the Government of Catalonia has provided funding to ME (2021 SGR01494). ME has also received funding from the Spanish Ministry of Science and Innovation MCIN/AEI/10.13039/501100011033/ERDF ‘A way to make Europe’ (PID2021-125282OB-I00), Cellex Foundation (CEL007) and “la Caixa” Foundation (LCF/PR/HR22/00732). M.E. is an ICREA Research Professor. GF received support by Fundacio La Marato de TV3 (ref 202131- 32). BP-M and JP is supported by Instituto de Salud Carlos III (ISCIII) (PI22/01892, PMP22/00054, PMP21/00107). EE is supported by Region Skane funds.

## Methods

### Generation of Visium Spatial Transcriptomics data from Formalin Fixed Paraffin Embedded fatal COVID-19 lung samples

First, the RNA integrity of the FFPE samples was assessed by extracting RNA from freshly collected tissue sections and evaluating the percentage of RNA fragments above 200 base pairs (DV200). Briefly, Tissue blocks were placed in the microtome (ThermoScientific HM340E) and trimmed to expose the tissue. 4 sections 10 µm thick were placed in a chilled Eppendorf tube and the RNA was extracted using a protocol from Qiagen (Rneasy FFPE Kit 73504), following extraction, the product was analyzed by TapeStation. Samples with DV200 ≥ 22% were selected for experiments.

Selected samples were placed in the microtome and sectioned 7 µm thick, each section was then placed in a water bath floating at 42 °C, sections were collected and mounted onto a 6.5 × 6.5 mm capture area of the Visium Spatial Gene Expression slide (2000233, 10X Genomics). Capture areas contain approximately 5000 barcoded spots, providing an average resolution of 1–10 cells. After sectioning, the slides were dried at 42°C for 3 hours. The slides were then placed inside a slide mailer, sealed with parafilm, and left overnight at Room temperature.

At the next day, the slides were deparaffinized by successive immersions in xylene and ethanol followed by H&E staining according to Demonstrated Protocol (CG000409, 10X Genomics). Brightfield images were taken using a 10X objective (Plan APO) on a Nikon Eclipse Ti2, images were stitched together using NIS-Elements software (Nikon) and exported as tiff files. After imaging, the glycerol and cover glass were carefully removed from the Visium slides by holding the slides in an 800 ml water beaker and letting the glycerol diffuse until the cover glass detached and density changes were no longer visible in the water. The slides were then dried at 37°C and Incubated for descrosslinking for 60 min.

Following decrosslinking step, over-night probe hybridization was performed, and libraries were prepared according to the Visium Spatial Gene Expression for FFPE User Guide (CG000407, 10X Genomics). Libraries were sent for sequencing in Macrogen Korea using 1 lane of HiSeq X 150PE (2x 150bp) per sample, applying 1% Phix. Sequencing was performed using the specific for FFPE following read protocol: read 1: 28 cycles; i7 index read: 10 cycles; i5 index read: 10 cycles; read 2: 50 cycles.

### Immunohistochemistry analysis

FFPE tissue sections were analyzed using standard IHC techniques. The primary antibodies used were anti-CD34 (clone QBEnd 10, Agilent Technologies, Santa Clara, CA, USA), anti-CD68 (clone KP1, Agilent Technologies, Santa Clara, CA, USA) and anti- CK7 (clone OV-TL 12/30, Agilent Technologies, Santa Clara, CA, USA). Immunostaining was performed automatically using a DAKO Autostainer Link 48 machine (Agilent Technologies, Santa Clara, CA, USA). Anti-CD34 was positive in endothelial cells, anti- CD68 was expressed in the cytoplasm of intraalveolar macrophages and CK7 was used as a marker for pneumocytes.

### Computational analysis

#### Visium Spatial Gene Expression libraries mapping

Visium Spatial Gene Expression libraries for Formalin Fixed Paraffin Embedded (FFPE) tissue samples were analyzed with spaceranger count pipeline using Space Ranger version 2.0.0 from 10x Genomics. First, a manual fiducial alignment and tissue boundary identification, including manual selection of spots covering tissue regions, were performed for each single library FFPE sample using Loupe Browser version 6.0 on the brightfield image. A probe set reference file compatible with FFPE workflow and human reference genome GRCh38 were downloaded from 10x Genomics and used to map Visium gene expression libraries.

#### Visium ST data preprocessing

We performed quality control (QC) steps including filtering of low-quality spots defined by a low number of detected genes with positive counts, low number of counts (library size) and high proportion of mitochondrial counts. These metrics were computed using scanpy (Wolf et al., 2018). As QC automatic filtering threshold, we utilized median absolute deviations (MAD) to identify outliers, as defined by differences in 5 MADs for number of detected genes and library size and 3 MADs for mitochondrial counts (including mitochondrial counts exceeding 8%) per tissue slide (Heumos et al., 2023).

We next applied normalization to the raw counts by scaling the counts followed by the shifted logarithm transformation to stabilize variance in gene expression between cells. To filter out uninformative genes with mostly zero counts, we performed feature selection using deviance to select informative genes (Heumos et al., 2023) using scry R package and selecting the top 6,000 highly deviant genes, as inspired by the preprocessing workflow utilized by the Human Lung Cell Atlas (Sikkema et al., 2023).

Finally, we performed dimensionality reduction using principal component analysis (PCA), t-distributed stochastic neighbor embedding (t-SNE) and uniform manifold approximation and projection (UMAP) with scanpy default parameters to reduce data complexity and for visualization purposes. To identify cellular structure, we cluster cells applying the Leiden algorithm to the previously computed neighborhood graph using different resolution parameters (0.25, 0.5, 1) with scanpy. Lastly, individual sample objects were joined into a single object using the anndata concat() function. After the concatenation, we re-normalized raw counts on the joined object using global scaling by the total counts per barcode and applying the shifted logarithm transformation, followed by feature selection using deviance to select the top 6,000 highly deviant genes, dimensionality reduction and clustering as previously described. A total of 91,068 spots, including 77,580 spots from fatal COVID-19 samples and 13,488 spots from control samples were profiled after QC.

#### Human Lung Cell Atlas (HLCA) reference processing

We leveraged the HLCA (Sikkema et al., 2023) single-cell RNA sequencing (scRNA-seq) reference dataset and consensus cell type annotations for spatial mapping and annotation. To this end, we downloaded and processed the HLCA core anndata object, selected lung parenchyma tissue cell types with at least 150 total cells at the finest level of annotation for a more robust and reliable reference model training, including a total of 333,011 cells in the filtered dataset. The mitochondrial genes were removed for spatial mapping.

#### Spatial mapping of cell types with cell2location

Both, our joined Visium ST and the filtered HLCA reference datasets, were subset to the same gene set as baseline for the mapping between single cell and spatial data, using default parameters to select a total number of 5,850 ENSEMBL gene identifiers. First, a reference model was fitted to estimate the reference cell type signature derived from the HLCA scRNA-seq data with cell2location (Kleshchevnikov et al., 2022) and using the finest level of cell type annotation reported. Cell2location uses a Negative Binomial regression model to estimate signatures, while accounting for batch effect and covariates. Hence, we included the following variables from the HLCA core object as covariates in our model: “assay”, “donor_id”, “tissue_sampling_method” and “tissue_dissociation_protocol”. To train the regression model we used default parameters to perform training on all cells in the dataset. A maximum number of 250 epochs were sufficient to achieve convergence.

For the subsequent spatial mapping, cell2location requires two user-provided hyperparameters based on the tissue and experiment QC, including expected number of cells per spot, that we set to 20, and regularization parameter of within slide or batch variation in RNA detection sensitivity, set to 20 (default), as previously described for the profiling of human lung tissue with Visium ST (Madissoon et al., 2023). The model was trained using full data until convergence with 40,000 iterations and loss function (ELBO) was used. Reconstruction accuracy plots were inspected to assess model quality. Cell abundance mapped to spatial coordinates was derived using the 5% quantile of the posterior distribution. To ease visualization of cell type abundances, the most abundant cell type per spot, including aggregation by cell type lineage, were represented. Marker gene selection was performed for every cell type by ranking genes using scanpy tool rank_genes_groups() function with default parameters using the t-test and computing a hierarchical clustering based on gene expression values for visualization with scaled expression for easier identification of differences. Additionally, cell-type specific expression of every gene at every spatial location was computed and used as input for cell-cell communication analysis with NCEM and for inferring intracellular signaling networks in peribronchial fibroblasts.

Lastly, we used non-negative matrix factorization (NMF) on cell2location mapping results to identify spatial co-occurrence of cell types. NMF was trained for a range of factors, selecting 8 factors for cellular compartments identification and visualization per condition with NMF factor loadings being represented.

#### Differential analysis of cell populations

To evaluate how cell populations changer across the studied biological conditions, we used scCODA model (Büttner et al., 2021) that employs a Bayesian model to perform compositional data analysis on the estimated cell-type abundances. The scCODA model determine statistically credible effects. We set the cutoff between credible and non-credible effects on a false discovery rate level (FDR) < 0.05. Estimated cell type abundances were used, including abundance aggregation by cell type lineage, and compositional data visualization was performed using stacked barplots and boxplots. To find a reference cell type that preserves changes in relative abundance across samples we used automatic reference cell type estimation, and Migratory dendritic cells (DCs) were deemed as reference category. Differences between conditions were computed using control samples as control group for control vs acute DAD and control vs proliferative DAD comparisons, whereas acute DAD samples were used as control group for acute DAD vs proliferative DAD comparisons.

#### Analysis of the spatial arrangement of cell types

A comparison of tissue architecture across conditions was performed leveraging novel statistical and computational approaches to compare cell spatial organization at the level of cell types and samples using GraphCompass (Ali et al., 2024). Samples are modelled as graphs of cells and cell-type specific graphs between conditions were compared with distances being computed using the portrait method and cell-type specific similarity scores were jointly visualized. Furthermore, comparisons between entire sample graphs for each cell type and condition were performed using filtration curves as previously described (Ali et al., 2024). For computing spatial graphs, we used default parameters for Visium ST samples and defined the cluster key as the spot-wise most abundant cell-type.

#### Cell-cell communication (CCC) analysis

To analyze intercellular communication events, we looked for potential ligand-receptor interactions (LRI) on our Visium ST slides using LIANA+ (Dimitrov et al., 2022; Dimitrov et al., 2023). To assess the spatial co-occurrence of LRIs we used spatially-informed local (individual spot-level) bivariate similarity metrics, including spatially-weighted Cosine similarity and local Morańs R (Dimitrov et al., 2023). These metrics use weights based on the spatial connectivity between spots, defined as radial kernels using the inverse Euclidean distance. Since we are assessing LRIs, we only considered interactions in which ligands, receptors and their subunits were expressed in at least 5% of the spots.

Furthermore, local interactions were categorized according to their magnitude and sign, allowing identification of local categories, where interactions are further classified into high-high (both variables are highly expressed) or high-low (one variable is highly and the other lowly expressed) interactions. Additionally, we used spot label permutations (N=100) to generate a Null distribution and to compute empirical local p-values to assess statistical significance of local metrics.

In addition to the local bivariate scores and to obtain “global” summaries of the local interaction results, we obtained global scores for each pair of variables, including Global mean (average) Cosine similarity and Global Morańs R (Dimitrov et al., 2023), to identify pairs of variables that cluster together or apart and to select the best candidates for visualization and downstream analysis.

Beyond LRIs, we applied the same approach to assess spatial relationships between transcription factors (TFs) activity (see below) and cell type abundances. To make the distributions comparable, we z-scaled TF activities and cell type abundances using zero-inflated minmax.

To identify coordinated cell-cell communication signatures, we used NMF on the local LRI scores. The heuristic elbow procedure selected three factors as the optimal component number. Average NMF factor scores per tissue slide clustered samples according to disease status and were visualized using a heatmap representation and hierarchical clustering using Euclidean distances and Ward’s method.

Pathway enrichment analysis of LRI loadings was performed to biologically characterize the three identified factors. To this end, pathway annotations from the PROGENy resource (Schubert M. et al., 2018) were converted into ligand-receptor sets as previously described (Dimitrov et al., 2023), assigning LRIs to specific pathways. Next, we performed enrichment using multivariate linear regression with decoupleR (Badia-i-Mompel et al., 2022). For LRI analysis we used LIANA+ consensus resource.

#### Transcription factor activity analysis

To infer TF activity based on prior knowledge, we used CollecTRI resource (Müller-Dott et al., 2023) containing a curated collection of TFs and their transcriptional targets with interactions weighted by their regulation mode (activation or inhibition). To estimate TF enrichment scores, we run a multivariate regression model using decoupleR for each spot and each TF, and a linear model was fitted to predict gene expression based on the interaction weights. The obtained t-value of the slope is the score, indicating activation or inactivation of the TF if positive or negative, respectively.

TFs enriched in each condition were identified using decoupleR rank_sources_groups() function with a t-test that overestimates variance of each group. The top 10 TFs per condition were extracted and represented in a heatmap using Z-scaled TF enrichment scores by standardizing scores to a comparable scale between 0 and 1, meaning for each TF, subtract the minimum and divide each by its maximum. Moreover, unstandardized TF enrichment scores were represented on top of illustrative Visium ST slides.

#### Learning spatial relationships between LRIs and TFs activity with cell type abundance

To learn spatial dependencies between local LRIs, TFs activity and cell types abundance, we used MISTy (Tanevski et al., 2022), an explainable multi-modelling approach, as previously described (Dimitrov et al., 2023). We selected the top 25 local ligand-receptor loadings from Factor 3 and the enrichment scores of the top 10 TFs per condition, to jointly modelled in a spatially informed manner the estimated cell types abundances in each spot. We specifically utilize a linear model, since the coefficients’ t-values (predictor importances), as calculated by ordinary least squares (OLS), are signed and comparable. Importantly, we bypassed predicting the intraview (intra spot), and for each target (cell type) we assessed not only the predictor importance, but how well LRIs and TFs explained cell type abundance using the joined multi-view R^2^ (goodness of fit) and evaluated their relative contribution to the joint predictive performance.

#### Intracellular signaling network in peribronchial fibroblasts

We inferred intracellular signaling networks from prior knowledge by linking identified LRIs and TF activity scores. LIANA+ approach (Dimitrov et al., 2023) considers the direction of deregulation, including activation and inhibition of receptors and TFs, and the sign and direction of edges (activating or inhibiting) from prior knowledge of protein-protein interactions and of TFs and their targets obtained from Omnipath (Turei et al., 2021). Using CORNETO, a putative causal network connecting LRIs to TFs was inferred for peribronchial fibroblasts. Hence, we used the coefficientś t-values of the predictive linear model for peribronchial fibroblasts abundance as input (LRIs) and output (TFs) nodes of the intracellular signaling network. To this end, we computed the median of the t-values across samples and ranked them by absolute median value. For LRIs, the receptors were selected and when the same receptor was involved in multiple LRIs, the largest absolute median t-value was kept for that receptor and used as input node. Additionally, we obtained a prior knowledge network (PKN) with signed protein-protein interactions from Omnipath and used peribronchial fibroblasts specific gene expression computed with cell2location to calculate gene expression proportions within our target cell type, using those to generate weights for the nodes in the PKN.

#### Analysis of intercellular dependencies as a function of spot composition

Since cell-cell communications events are not limited to LRIs, we further assessed spot/niche composition effects on all genes using NCEM (Fischer et al., 2023). To this end, we utilized cell-type specific gene expression computed with cell2location. Hence, NCEM models expression variation within cell types across spots as function of the inferred spot composition. To focus the analysis on biologically relevant genes, we selected gene sets described in the WikiPathways database from the Molecular Signature Database (MSigDB) using decoupleR. We filtered out lineage specific marker genes computed using rank_genes_groups_df() function in scanpy using adjusted p-value < 0.05 and minimum log fold change > 2. Finally, 1614 genes were shared within our dataset. Additionally, 22 cell types were considered for NCEM analysis, including cell types with credible differential abundances computed by scCODA and the most abundant cell types.

Type coupling analysis was performed on the filtered dataset to compute sender and receiver effects based on a Wald test on the parameters learned by the linear NCEM model, using the full dataset and optimized with OLS. We further dissected these couplings based on gene-wise effects of particular interactions, including effects of all senders on one receiver (receiver effect analysis), reporting dependencies with at least 500 differentially expressed with q-value < 0.05 and absolute log fold change > 0.8. Finally, a sender similarity analysis was performed to characterize sender profiles on CD4 T cells across conditions.

#### Spatio-temporal trajectories analysis

To characterize AT2-AT1 differentiation process, we leveraged stLearn tool (Pham et al. 2023) that employs a spatial graph-based method named pseudo-time-space (PSTS) that combines spatial and imaging information with gene expression to map spatial changes cell states, modelling and reconstructing their spatio-temporal trajectories. To accurately identify transition genes positively or negatively correlated with the predicted trajectory for AT2-AT1 differentiation, we reported genes with a Spearman correlation <- 0.4 or > 0.4. Furthermore, we only selected spots enriched in AT1 and AT2 cells, where these cell types represented at least 70% of the maximum abundance of the inferred spot composition.

---

**Supplementary Figure 1:**
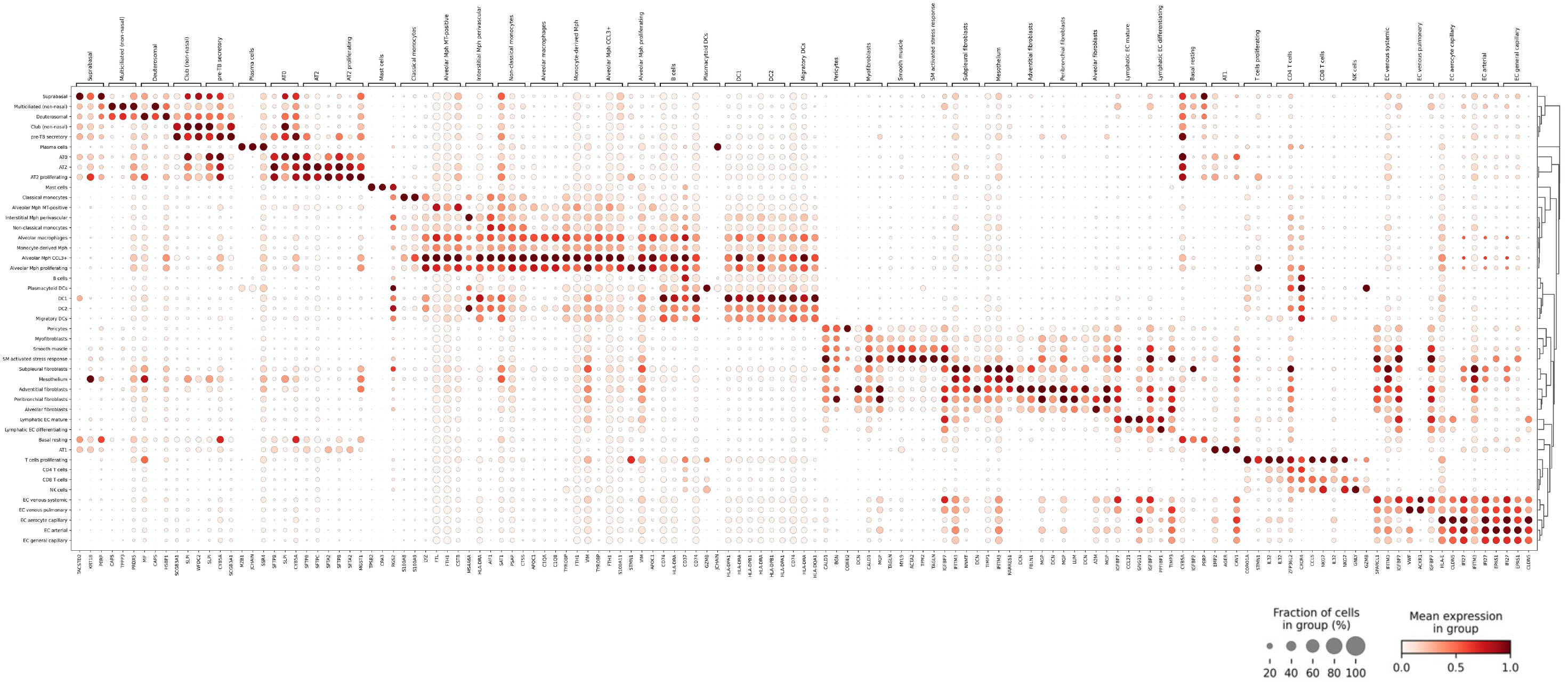
Marker gene expression of reference signatures derived from the HLCA core for all 45 cell types identified. Gene expression was normalized such that the maximum group expression of cells for each marker was set to 1. Dot size indicates the fraction of cells in group.

**Supplementary Figure 2:**
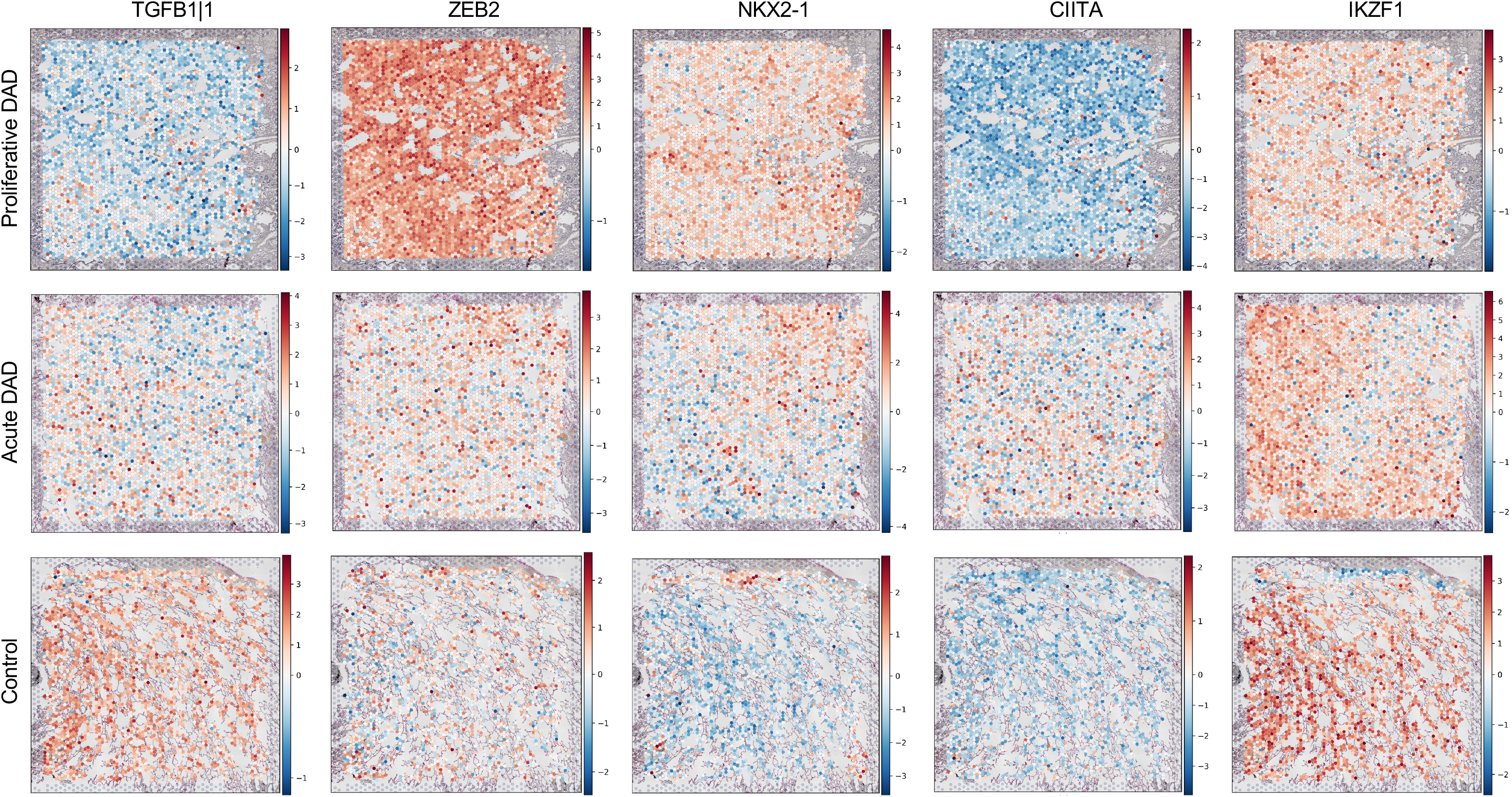
Transcription factors activity across disease progression. Enrichment scores of selected transcription factors across disease progression in illustrative Visium ST lung samples are shown.

**Supplementary Figure 3:**
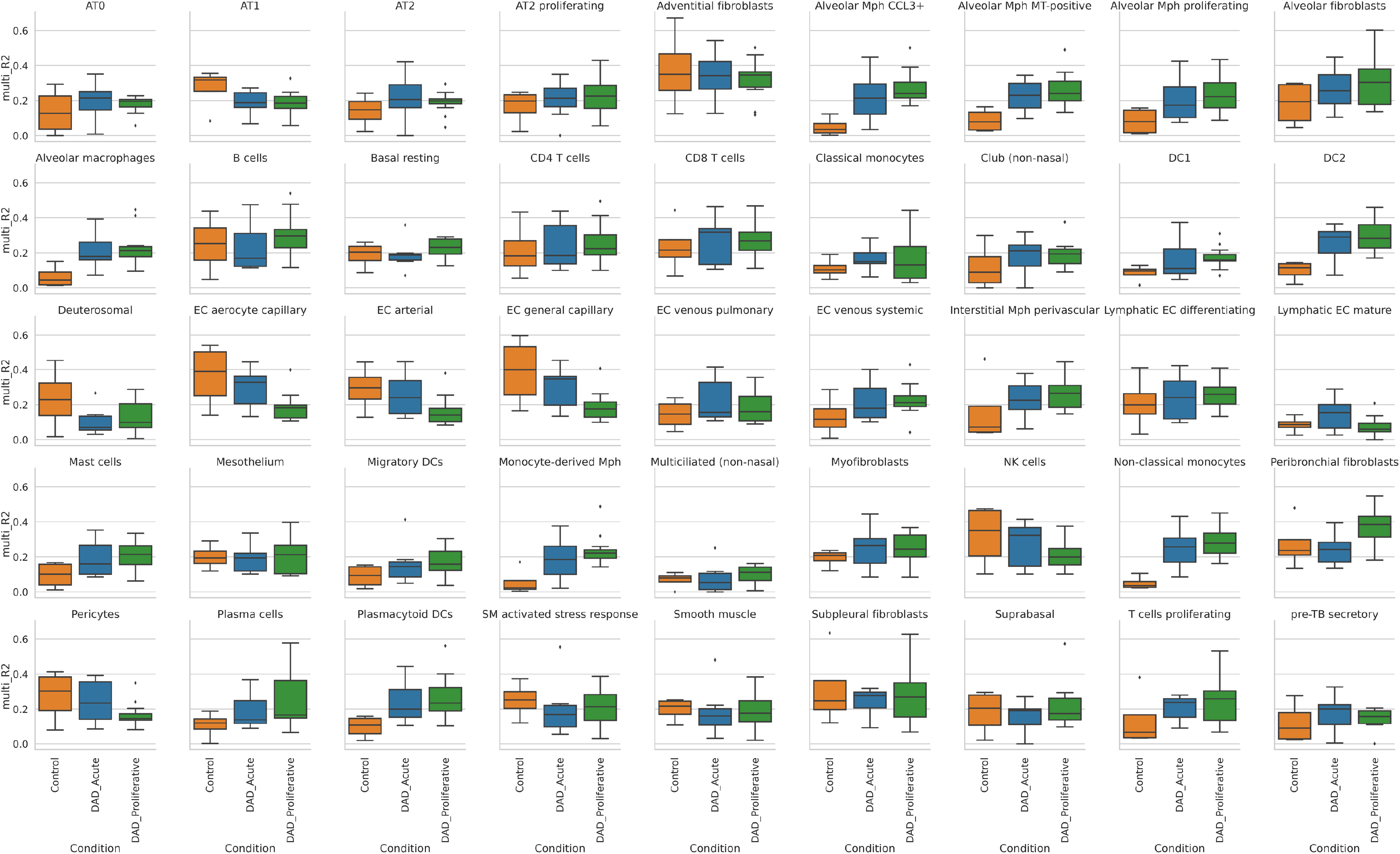
Variance of cell type abundance explained by the predictive multi-view model. The joint LRI and TF multi-view R^2^ contributing to explain cell type abundance variance across disease porgression is shown.

**Supplementary Figure 4:**
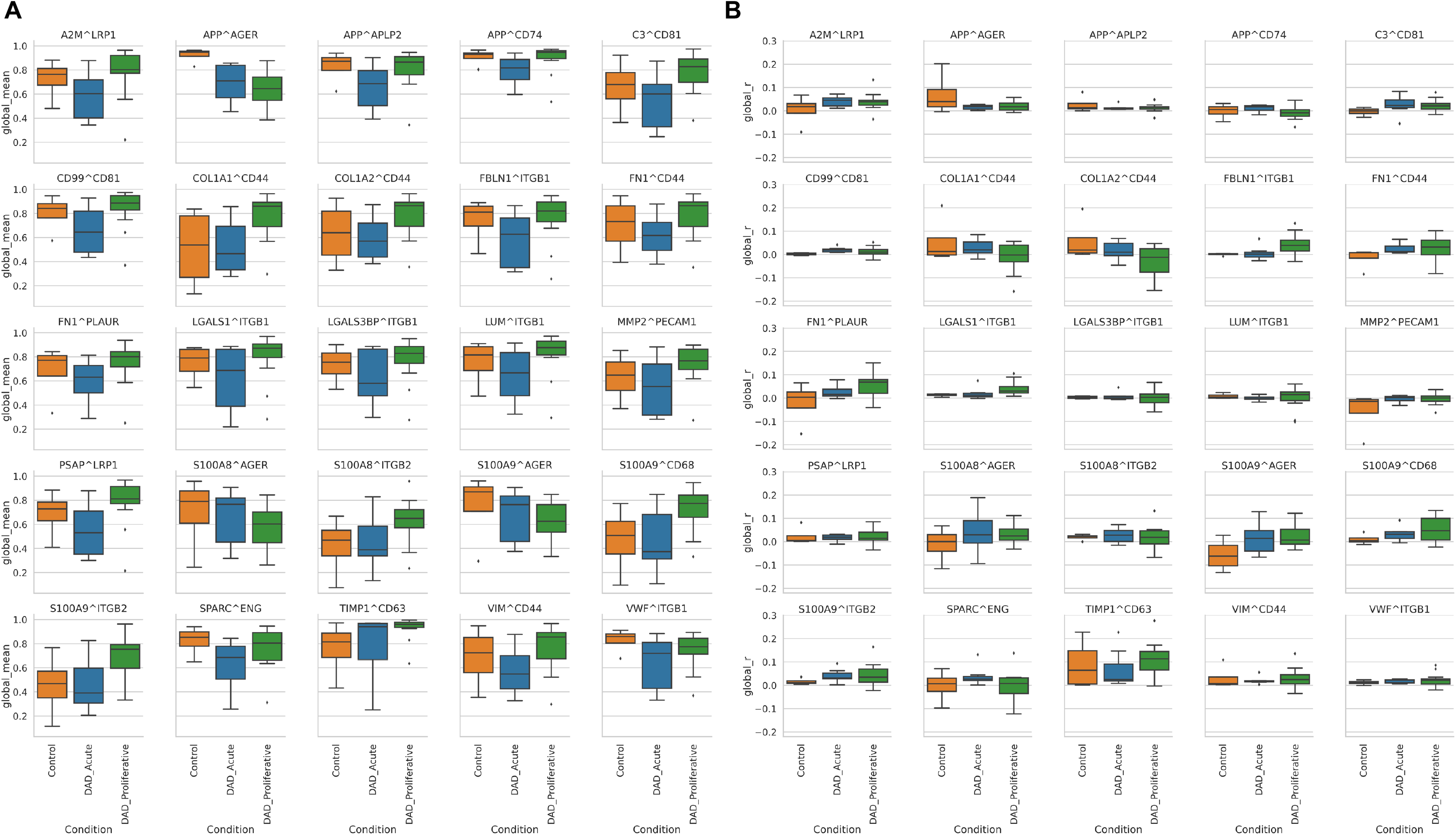
Local spatial relationships of ligand-receptor interactions across disease progression. **A)** Global mean Cosine similarity and **B)** Global Morańs R for the top 25 LRI defining DAD associated Factor 3 across disease progression are shown.

**Supplementary Figure 5:**
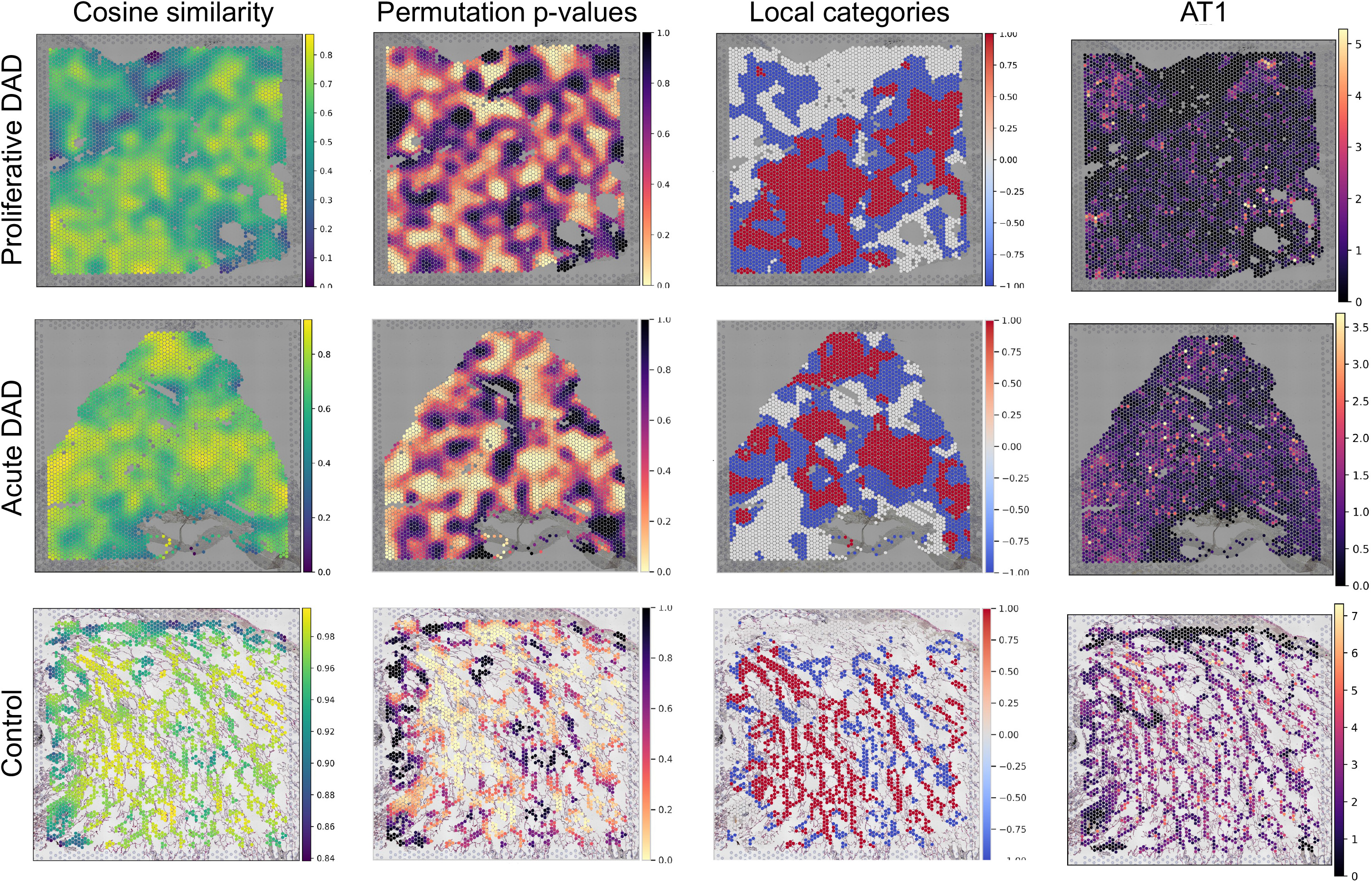
Local spatial relationship of APP^AGER ligand-receptor interaction. Cosine similarity, permutation p-values, local categories and AT1 cell type abundance on illustrative Visium ST lung samples across disease progression are shown. High-high interactions (red) and high-low or low-high interactions (blue) are depicted in local categories plots.

**Supplementary Figure 6:**
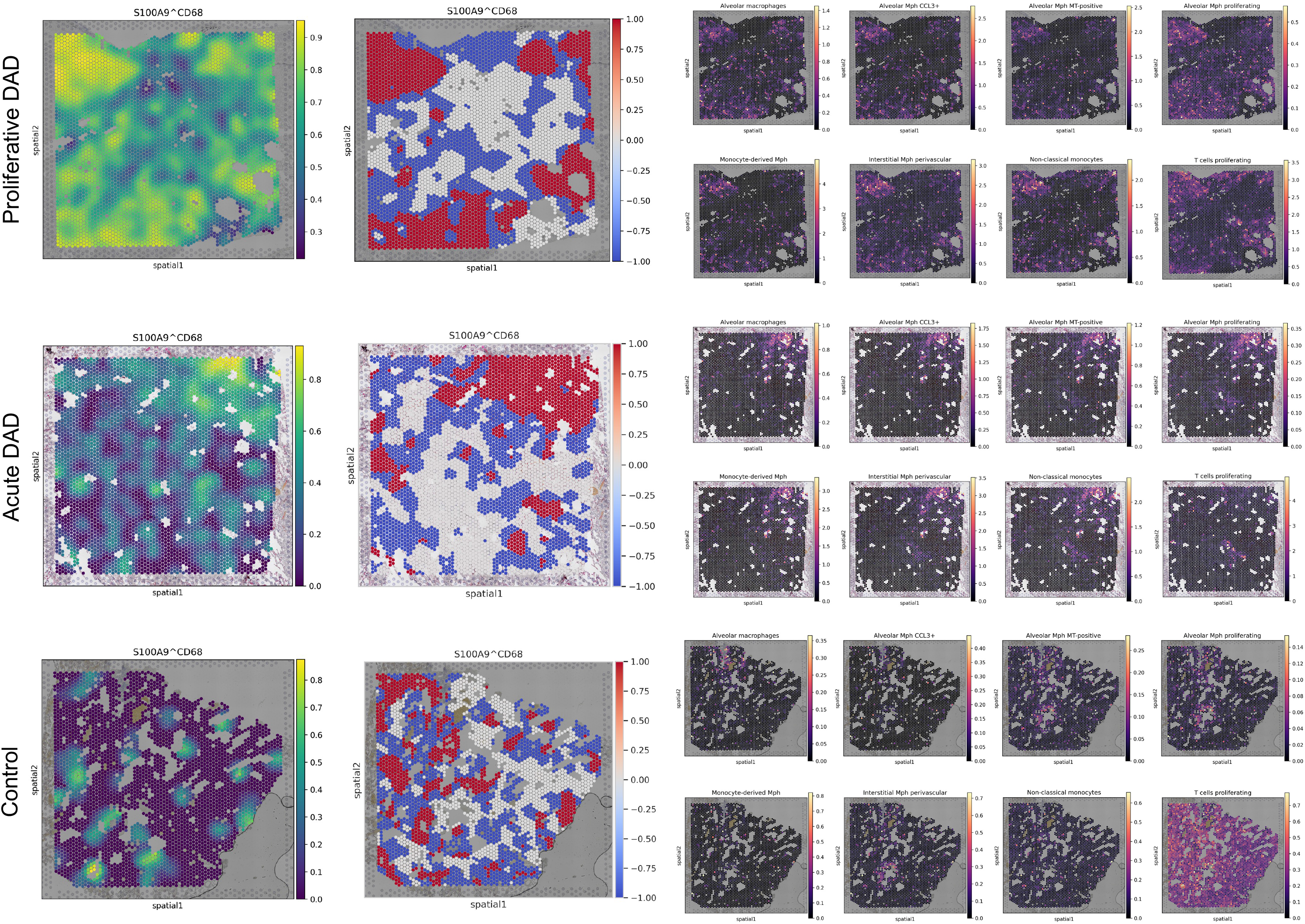
Local spatial relationship of S100A9^CD68 ligand-receptor interaction. Cosine similarity, local categories, myeloid and T cells proliferating cell type abundance on illustrative Visium ST lung samples across disease progression are shown. High-high interactions (red) and highlow or low-high interactions (blue) are depicted in local categories plots.

**Supplementary Figure 7:**
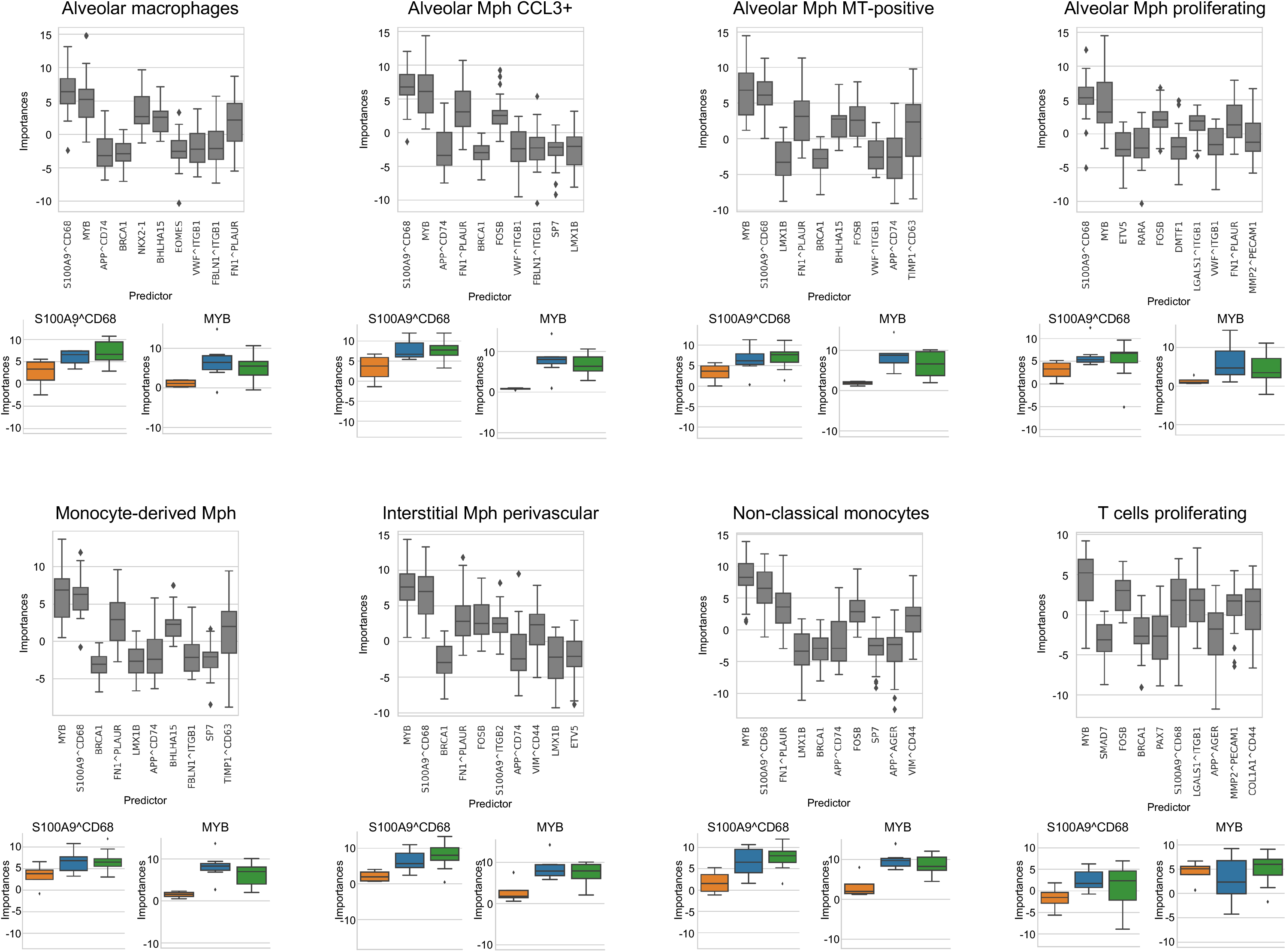
S100A9^CD68 and MYB predictive performance for myeloid and T cells proliferating cell types abundance. The top 10 predictors of myeloid cell types and T cells proliferating abundances are shown. Additionally, S100A9^CD68 LRI and MYB TF predictor importances for the same cell types across disease progression are shown.

**Supplementary Figure 8:**
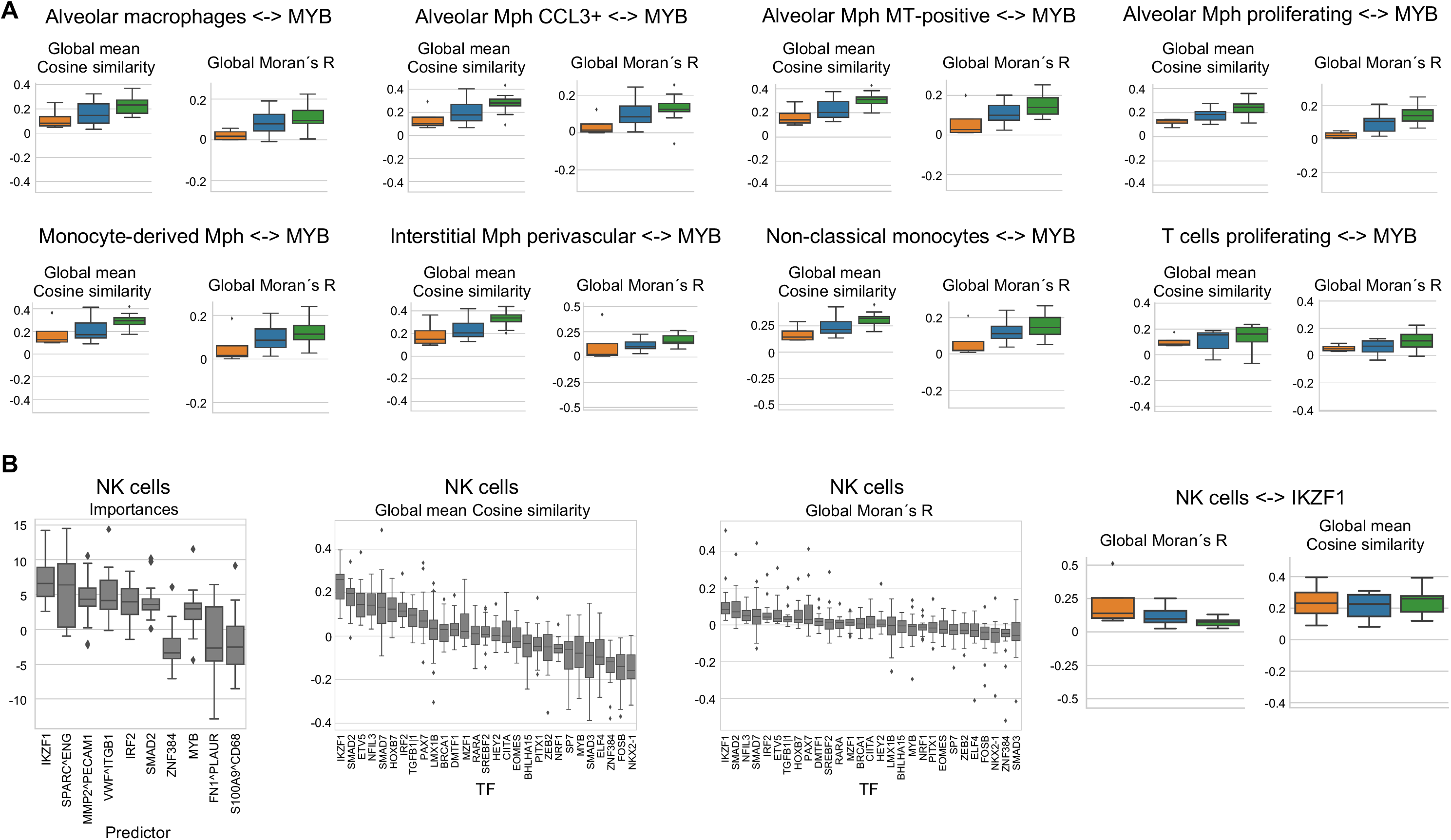
Local spatial relationship of MYB transcription factor. **A)** Global mean Cosine similarity and Global Morańs R for myeloid cell types and T cells proliferating with MYB are shown across disease progression. B) Top 10 predictors of NK cells, Global mean Cosine similarity and Global Morańs R for spatial co-clustering of NK cells with the top 10 TFs enrichment scores per condition, and NK cells spatial co-occurrence across disease progression are shown.

